## Supplemental material for "Predicting individual differences of fear and cognitive learning and extinction"

#### Supplemental Methods

##### Preprocessing (fmriprep boilerplate)

Results included in this manuscript come from preprocessing performed using fMRIPrep 20.1.1 (Esteban, Markiewicz, et al. (2018); Esteban, Blair, et al. (2018); RRID:SCR_016216), which is based on Nipype 1.5.0 (Gorgolewski et al. (2011); Gorgolewski et al. (2018); RRID:SCR_002502).

Anatomical data preprocessing

A total of 1 T1-weighted (T1w) images were found within the input BIDS dataset.The T1-weighted (T1w) image was corrected for intensity non-uniformity (INU) with N4BiasFieldCorrection (Tustison et al. 2010), distributed with ANTs 2.2.0 (Avants et al. 2008, RRID:SCR_004757), and used as T1w-reference throughout the workflow. The T1w-reference was then skull-stripped with a Nipype implementation of the antsBrainExtraction.sh workflow (from ANTs), using OASIS30ANTs as target template. Brain tissue segmentation of cerebrospinal fluid (CSF), white-matter (WM) and gray-matter (GM) was performed on the brain-extracted T1w using fast (FSL 5.0.9, RRID:SCR_002823, Zhang, Brady, and Smith 2001). Brain surfaces were reconstructed using recon-all (FreeSurfer 6.0.1, RRID:SCR_001847, Dale, Fischl, and Sereno 1999), and the brain mask estimated previously was refined with a custom variation of the method to reconcile ANTs-derived and FreeSurfer-derived segmentations of the cortical gray-matter of Mindboggle (RRID:SCR_002438, Klein et al. 2017). Volume-based spatial normalization to two standard spaces (MNI152NLin6Asym, MNI152NLin2009cAsym) was performed through nonlinear registration with antsRegistration (ANTs 2.2.0), using brain-extracted versions of both T1w reference and the T1w template. The following templates were selected for spatial normalization: FSL's MNI ICBM 152 non-linear 6th Generation Asymmetric Average Brain Stereotaxic Registration Model [Evans et al. (2012), RRID:SCR_002823; TemplateFlow ID: MNI152NLin6Asym], ICBM 152 Nonlinear Asymmetrical template version 2009c [Fonov et al. (2009), RRID:SCR_008796; TemplateFlow ID: MNI152NLin2009cAsym],

Functional data preprocessing

For each of the 1 BOLD runs found per subject (across all tasks and sessions), the following preprocessing was performed. First, a reference volume and its skull-stripped version were generated using a custom methodology of fMRIPrep. Head-motion parameters with respect to the BOLD reference (transformation matrices, and six corresponding rotation and translation parameters) are estimated before any spatiotemporal filtering using mcflirt (FSL 5.0.9, Jenkinson et al. 2002). BOLD runs were slice-time corrected using 3dTshift from AFNI 20160207 (Cox and Hyde 1997, RRID:SCR_005927). Susceptibility distortion correction (SDC) was omitted. The BOLD reference was then co-registered to the T1w reference using bbregister (FreeSurfer) which implements boundary-based registration (Greve and Fischl 2009). Co-registration was configured with six degrees of freedom. The BOLD time-series were resampled onto the following surfaces (FreeSurfer reconstruction nomenclature): fsaverage. The BOLD time-series (including slice-timing correction when applied) were resampled onto their original, native space by applying the transforms to correct for head-motion. These resampled BOLD time-series will be referred to as preprocessed BOLD in original space, or just preprocessed BOLD. The BOLD time-series were resampled into standard space, generating a preprocessed BOLD run in MNI152NLin6Asym space. First, a reference volume and its skull-stripped version were generated using a custom methodology of fMRIPrep. Grayordinates files (Glasser et al. 2013) containing 91k samples were also generated using the highest-resolution fsaverage as intermediate standardized surface space. Automatic removal of motion artifacts using independent component analysis (ICA-AROMA, Pruim et al. 2015) was performed on the preprocessed BOLD on MNI space time-series after removal of non-steady state volumes and spatial smoothing with an isotropic, Gaussian kernel of 6mm FWHM (full-width half-maximum). Corresponding "non-aggresively" denoised runs were produced after such smoothing. Additionally, the "aggressive" noise-regressors were collected and placed in the corresponding confounds file. Several confounding time-series were calculated based on the preprocessed BOLD: framewise displacement (FD), DVARS and three region-wise global signals. FD was computed using two formulations following Power (absolute sum of relative motions, Power et al. (2014)) and Jenkinson (relative root mean square displacement between affines, Jenkinson et al. (2002)). FD and DVARS are calculated for each functional run, both using their implementations in Nipype (following the definitions by Power et al. 2014). The three global signals are extracted within the CSF, the WM, and the whole-brain masks. Additionally, a set of physiological regressors were extracted to allow for component-based noise correction (CompCor, Behzadi et al. 2007). Principal components are estimated after high-pass filtering the preprocessed BOLD time-series (using a discrete cosine filter with 128s cut-off) for the two CompCor variants: temporal (tCompCor) and anatomical (aCompCor). tCompCor components are then calculated from the top 5% variable voxels within a mask covering the subcortical regions. This subcortical mask is obtained by heavily eroding the brain mask, which ensures it does not include cortical GM regions. For aCompCor, components are calculated within the intersection of the aforementioned mask and the union of CSF and WM masks calculated in T1w space, after their projection to the native space of each functional run (using the inverse BOLD-to-T1w transformation). Components are also calculated separately within the WM and CSF masks. For each CompCor decomposition, the k components with the largest singular values are retained, such that the retained components' time series are sufficient to explain 50 percent of variance across the nuisance mask (CSF, WM, combined, or temporal). The remaining components are dropped from consideration. The head-motion estimates calculated in the correction step were also placed within the corresponding confounds file. The confound time series derived from head motion estimates and global signals were expanded with the inclusion of temporal derivatives and quadratic terms for each (Satterthwaite et al. 2013). Frames that exceeded a threshold of 0.5 mm FD or 1.5 standardised DVARS were annotated as motion outliers. All resamplings can be performed with a single interpolation step by composing all the pertinent transformations (i.e. head-motion transform matrices, susceptibility distortion correction when available, and co-registrations to anatomical and output spaces). Gridded (volumetric) resamplings were performed using antsApplyTransforms (ANTs), configured with Lanczos interpolation to minimize the smoothing effects of other kernels (Lanczos 1964). Non-gridded (surface) resamplings were performed using mri_vol2surf (FreeSurfer).

Many internal operations of *fMRIPrep* use *Nilearn* 0.6.2 (Abraham et al. 2014, RRID:SCR_001362), mostly within the functional processing workflow. For more details of the pipeline, see the section corresponding to workflows in *fMRIPrep*'s documentation.

**Spectral Dynamic Causal Modelling**

Dynamic causal modelling (DCM) is a Bayesian framework that infers the directed (causal) connectivity among the neuronal systems – referred to as effective connectivity. We recently proposed a new DCM for resting state fMRI – based upon a deterministic model that generates predicted cross spectra – referred to as spectral DCM. In order to model resting state activity – in the absence of external stimuli – we will have to add a stochastic component, i.e. neural fluctuations, to the classical DCM based on ordinary differential equations. Mathematically, we can express the formulation of the stochastic generative model using a set of two equations. First is the neuronal state equation, namely

$$\dot{x}(t) = f(x(t), u(t), \theta) + v(t), \tag{S1}$$

and second is the observation equation, which is a static nonlinear mapping from the hidden physiological states in (1) to the observed BOLD activity and is written as:

$$y(t) = h(x(t), \varphi) + e(t), \tag{S2}$$

where $\dot{x}(\mathrm{t})$ is the rate of change of the neuronal states $x(t)$, $\theta$ are unknown parameters (i.e. the effective connectivity) and $v(t)$ (resp. $e(t)$) is the stochastic process – called the state noise (resp. the measurement or observation noise) – modelling the random neuronal fluctuations that drive the resting state activity. In the observation equations, $\varphi$ are the unknown parameters of the (haemodynamic) observation function and $u(t)$ represents any exogenous (or experimental) inputs that drive the hidden states – that are usually absent in resting state designs [1]. Spectral DCM furnishes a constrained inversion of the stochastic model by parameterising the neuronal fluctuations $v(t)$. Spectral DCM simplifies the generative model by replacing the original timeseries with their second-order statistics (i.e., cross spectra). This means, instead of estimating time varying hidden states, we are estimating their covariance which is time invariant. Then we simply need to estimate the covariance of the random fluctuations; where a scale free (power law) form for the state noise (resp. observation noise) is used – motivated from previous work on neuronal activity [2–4] – as follows:

$$g_v(\omega, \theta) = \alpha_v \omega^{-\beta_v}$$

$$g_e(\omega, \theta) = \alpha_e \omega^{-\beta_e} \tag{S3}$$

Here, $\{\alpha, \beta\} \subset \theta$ are the parameters controlling the amplitudes and exponents of the spectral density of the neural fluctuations. The parameterisation of endogenous fluctuations means that the states are no longer probabilistic; hence the inversion scheme is significantly simpler, requiring estimation of only the parameters (and hyperparameters) of the model.

We used standard Bayesian model inversion to infer the parameters of the model in (1), (2) and (3), from the observed signal $y(t)$. The description of the Bayesian model inversion procedures based on variational Laplace can be found elsewhere for the interested readers [5–7].

**Parametric Empirical Bayes**

Empirical Bayes refers to the Bayesian inversion or fitting of hierarchical models. In hierarchical models, constraints on the posterior density over model parameters at any given level are provided by the level above. These constraints are called empirical priors because they are informed by empirical data. We recently introduced a second-level or between-subjects model over parameters, which represents how individual (within-subject) connections derive from the subjects' group membership[8] – based on parametric empirical Bayes (PEB). This approach calls on Bayesian Model Reduction (BMR) to finesse the inversion of multiple models of a single dataset or a single (hierarchical) model of multiple datasets. BMR allows one to compute posterior densities over model parameters, under new prior densities, without explicitly inverting the model again. For example, one can invert a DCM for each subject in a group and then evaluate the posterior density over group effects, using the posterior densities over parameters from the single subject inversion. This may improve subject-specific parameter estimates, by using group-level estimates to rescue individual DCM from local optima. Mathematically, for DCM studies with $N$ subjects and $M$ parameters per DCM, we have a hierarchical model, where the responses of the $i$-th subject and the distribution of the parameters over subjects can be modelled as:

$$y_i = \Gamma_i^{(1)}\left(\theta^{(1)}\right) + \varepsilon_i^{(1)} \qquad \text{(S4)}$$
$$\theta^{(1)} = \Gamma^{(2)}\left(\theta^{(2)}\right) + \varepsilon^{(2)}$$
$$\theta^{(2)} = \eta + \varepsilon^{(3)}$$

where, $y_i$ is the BOLD time series from *i-th* subject *and* $\Gamma_i^{(1)}$ is a nonlinear mapping from the parameters of a model to the predicted response $y$, for example, as shown in Eq. S1 above. $\varepsilon_i^{(1)}$ is independent and identically distributed (i.i.d.) observation noise (equivalent to $e(t)$ in Eq. S2). In this hierarchical form, *empirical priors* encoding second (between-subject) level effects place constraints on subject-specific parameters. The second level would be a linear model where the random effects are parameterised in terms of their precision:

$$\Gamma^{(2)}\left(\theta^{(2)}\right) = (X \otimes W)\beta$$

where, $\beta \subset \theta$ are group means or effects encoded by a design matrix with between-subject $X$ and within-subject $W$parts. The between-subject part encodes differences among subjects or covariates such as age, while the within-subject part specifies mixtures of parameters that show random effects. We assume that the first column of the design matrix is a constant term, modelling group means and subsequent columns encode group differences or covariates such as age.

**Functional connectivity metrics**

Pearson correlation (cor) captures linear dependencies between two signals. Correlation values are bounded between -1 and 1. Positive correlation values indicate that as one signal increases in strength so does the other. Negative values, on the other hand, imply that the signals move in opposite directions. Importantly, correlation values close to 0 simply suggest that there is little or no linear relationship; it does not mean that a relationship does not exist.

Empirical covariances and empirical Pearson correlations have been found to be subpar statistical estimators of the “true” covariances that would be observed had we performed an infinite number of measurements. To circumvent this issue, a Ledoit-Wolf estimator was used[9], which corrects empirical covariances by shrinking their variability around their average:

$$cov_{shrunk} = (1 - \delta) * cov + \delta \frac{Tr(cov)}{p} Id$$

Cross-correlation (xcor) is related to Pearson correlation in that it also captures linear dependencies among two signals. However, it differs in that it calculates linear correlations between shifted versions of one signal with respect to another. Thus, cross-correlation produces a vector of similarity measures, as opposed to a single correlation value as in Pearson correlation. In our current approach, we extracted the maximum value of this vector, which may be different from Pearson correlation if two signals happen to be most strongly correlated with a nonzero lag.

Dynamic time warping (dtw) is a scale-invariant dissimilarity measure that quantifies how dissimilar two signals are. It warps the time axis in a non-linear fashion to provide the best alignment between the signals. DTW ranges from 0 to infinity, with values closer to 0 indicating good alignment between the two time series.

For our implementation of dtw, we used a symmetric recursion step pattern that can be normalised by the length of the time series, a Sakoe-Chiba window of size = 100 (enforcing a global constraint on the warping path[10]), and squared Euclidean distance as distance metric.

Euclidean distance (ED) is a dissimilarity measure that simply measures the geometric distance between any two points in Euclidean space. It bounds at 0 at the lower end, but is unbounded above. The closer the values are to 0 the greater the overlap between the two time series.

Manhattan distance (MD) is computed by taking the sum of the absolute differences between two time series. As with ED, it bounds at 0 at the lower end, and is unbounded above, with values closer to 0 indicating greater overlap.

Wasserstein distance (WD) is a dissimilarity metric that treats each signal as a probability distribution. WD can be seen as the minimum amount of work needed to convert one probability distribution into the other (i.e., the amount of distribution weight that needs to be moved times the distance).

Mutual information (MI) is a measure of mutual non-linear dependence between two signals. In essence, it quantifies how much information can be gathered from one signal by observing the behaviour of another. A value of 0, therefore, indicates that knowing one of the signals provides no knowledge of the other signal (i.e., they are independent). In contrast, as MI values increase, so does the reduction of uncertainty (i.e., there is a deterministic relationship between the signals).

Magnitude-square coherence (cohe) indicates how correlated the two signals are in the frequency-domain. Values are bounded between 0 (no coherence) and 1 (complete coherence). cohe outputs a value for each frequency, thus producing a vector. Commonly, the maximum value of this vector is taken to indicate the strongest coherence achievable.

Wavelet coherence (wcohe) measures the correlation between two time-series on the time-frequency plane by employing the wavelet power spectrum of the two signals (we used an analytic Morlet wavelet to compute coherence). wcohe is a useful metric to analyse non-stationary signals and has advantages over other correlation methods in that it looks at coherence coefficients at different scales.

For the calculation of the composite score, the absolute value of cor was used, and the values for the distance metrics (ED, MD, WD and dtw) were inverted, so that 0 reflected no association and 1 a perfect association. Next, we concatenated the different metric values by simple summation, producing a single score per ROI pair and per subject.

#### Psychophysiological model for SCR

For analysis of SCR data, we opted for a psychophysiological modelling approach, which is conceptually and mathematically related to the way we analyse our fMRI data. Time-series of observed SCR data are regarded as the output of a generative psychophysiological model with unknown input, and the input is estimated from the data. This approach makes use of the entire time series of data, rather than just selected features such as peaks and troughs. Compared to several traditional SCR analysis approaches, it is often more sensitive in recovering known effects of an experimental manipulation in controlled experiments[11,12] (a validation approach termed "experiment-based calibration"[13,14]).

The non-linear psychophysiological model (PsPM) for SCR is formulated in terms of linear dynamic systems (LDS) with time-dependent input[15]. The LDS model the periperal system, i.e. the conduction delay of the sudomotor nerve, diffusion of ACh to the sweat glands, the sweat gland response, and the evaporation of sweat. The time-dependent input models the neural input into the sudomotor system. The amplitude of the anticipatory, CS-related input is taken as an index of the current US (fear) memory.

*Neural system*

Neural inputs are modelled as Gaussian bumps with estimated amplitude $a$. For each experimental event:

$$u_{exp}(t) = a\exp\frac{-(t-\mu)^2}{2\sigma^2},$$

with fixed $\sigma = 0.3$ seconds. We defined fixed $\mu = t_{context}$ or $\mu = t_{US}$ for context- and US-related responses (corresponding to an immediate neural firing burst), and estimated $t_{CS} < \mu < t_{US}$ for CS-related responses (corresponding to a neural firing burst at some point between CS onset and US onset), where $t_{context}$, $t_{CS}$ and $t_{US}$ are the respective context, CS and US onset times.

Furthermore, we modelled spontaneous neural firing bursts $u_{sf}(t)$ (often termed non-specific SCR, or spontaneous fluctuations) during inter-trial-intervals in the same way, where we fixed $\sigma = 0.3$ seconds and estimated $\mu_{min} < \mu < \mu_{max}$. For each inter-trial-interval, $\mu_{min}$ corresponds to the last US onset plus 5 seconds, and $\mu_{max}$ to the next trial onset minus 2 seconds, thus ensuring that the modelled SF do not absorb variance caused by the experiment. During this interval, the number of modelled SF corresponded to 1 SF per 2 seconds.

Finally, we modelled residual baseline fluctuations between trials $u_{scl}(t)$ in a similar way, with fixed $\sigma_0 = 1.0$ and estimated $\mu_{min} < \mu < \mu_{max}$, where the limits are the same as for SF.

*Peripheral system*

The peripheral system was modelled as a sum of three LDS; each taking one component of the neural input. The LDS corresponding to experimental events and SF are of the same form but with different response functions, which were empirically determined in previous work. Both LDS were of the form:

$$\dddot{y} + \vartheta_1\ddot{y} + \vartheta_2\dot{y} + \vartheta_3 y - u(t-\vartheta_4) = 0;$$

where $y(t)$ are the SCR data, dot notation is used for time derivatives, and $u$ corresponds to $u_{exp}$ for the first LDS and to $u_{sf}$ for the second LDS. Parameters for the first LDS are $\hat{\vartheta}_1 = 1.342052$, $\hat{\vartheta}_2 = 1.411425$, $\hat{\vartheta}_3 = 0.122505$, $\hat{\vartheta}_4 = 1.533879$ (estimated previously from 1278 experiment-evoked SCRs from 64 individuals[16]); and for the second LDS $\hat{\vartheta}_1 = 2.1594$, $\hat{\vartheta}_2 = 3.9210$, $\hat{\vartheta}_3 = 0.9235$, $\hat{\vartheta}_4 =$

0 seconds (estimated previously from 1153 semi-automatically detected SF from 40 individuals[17]). Amplitude of LDS is rescaled such that a canonical Gaussian bump SN impulse with unit amplitude elicits an SCR with unit amplitude. The third LDS was simply $y - u_{scl}(t) = 0$.

*Model inversion*

The model is inverted with the VBA toolbox. Because the parameter space is too large to invert the model at once, inversion is done in a trial-by-trial approach: a number of trials, $T_{1..n}$ is inverted. The ensuing parameter estimates for $T_1$ are extracted, and the parameter estimates for $T_{2..n}$ are used as prior values for the next inversion for trials $T_{2..n+1}$ where the estimated hidden state values at the start of trial 2 are taken as starting values of the hidden states $x$. This is continued until all trials have been estimated. This scheme ensures that inversion is kept tractable, and at the same time overlapping responses are taken into account. We have previously shown that inverting 2 or 3 trials at the same time yields comparable retrodictive validity[12].

**Experimental paradigms (extended text)**

S1: The full reversal fear paradigm was carried out in the MRI scanner and consisted of four experimental phases spread over two days. In each phase, participants were presented, during each trial, with pictures of household appliances for 1 second (e.g., washing machine, coffee maker), which were used as conditioned stimuli (CS) for a total of 16 possible CS. Some CS were paired with a US (electric shock) delivered for 0.75 seconds, while some CS were not paired with a US. In addition, the CS were embedded in and preceded (for 2 seconds) by a video of a scene that served as a context for the CS. The experimental phases differed in the way the CS were associated with a US (given 4 possible types of CS-US associations, or CS types, across experimental phases; see details below), as well as in the type of context in which the CS were embedded. US expectancy ratings were collected at each trial using a 4-point Likert scale during 2.5 seconds before administration (or lack of administration) of the electric shock. Visual stimuli were presented using the Presentation software package (Neurobehavioral Systems, Berkeley, CA, USA). Electrical stimulation was delivered using a constant voltage stimulator (STM2000, BIOPAC Systems, Goleta, CA, USA) with electrodes attached to the hand. The intensity of the electrical stimulation was adjusted individually for each participant prior to the fear acquisition phase until the participants rated the sensation as unpleasant but not painful. A fixation cross was shown with a jittered duration (7 to 9 seconds) to serve as intertrial interval at the end of each trial. Each experimental phase had 128 trials in total.

The first day of the experiment consisted two phases which we will refer to as "fear acquisition" and "fear reversal", respectively. During the fear acquisition phase, half of the CS types (out of 4 possible types, with two different cues per type; hence, 8 cues) were associated with a US (50% reinforcement) while the other half were not. The presentation of CS was embedded in the context of a natural scene (out of 4 possible scenes), with CS contingency and context type being orthogonal from each other. The fear acquisition phase was immediately followed fear reversal. Here, participants were again presented with the same CS. However, half of the CS associated with a US during fear acquisition were no longer associated with a US (corresponding to CS type named CS+-), while the other half remained associated with a US (the CS++ type). Similarly, half of the CS- cues became associated with a US (the CS-+ type), while the other half remained unassociated with a US (the CS-- type). Both CS++ and CS-+ cues were reinforced in 50% of the trials. The second day of the experiment consisted of two experimental phases, which we will refer to as fear extinction and fear renewal, respectively. During extinction and renewal, none of the CS types were associated with a US, but extinction and renewal differed in terms of the context videos in which the CS were embedded (see details below). Therefore, regarding the CS, the contingency of CS types was manipulated throughout the experimental phases in a way specific to each of the four types of CS. To summarize, CS++ cues refer to CS that are associated with a US in both the fear acquisition and the fear reversal phases; CS-+ refers to cues that are not associated with a US during fear acquisition but are associated with a US during reversal; CS+- refers to cues associated with a US during fear acquisition that are not associated with a US in fear reversal; CS-- refers to cues that are not associated with a US in fear acquisition and fear reversal. Although none of the CS types were associated with a US on the second day of the experiment, we refer to the CS types by their contingency changes between the first two experimental phases, since one of the goals of the paradigm was to understand how the contingency changes made between the fear acquisition and fear reversal phases influence the neural representations of these cues during fear extinction and fear renewal.

In all phases, context videos and CS types were presented in different pseudorandom orders such as to be orthogonal from each other. A set of 4 possible videos were shown during fear acquisition, a new set of 4 videos during fear reversal, and a new set of 8 videos during fear extinction. During fear renewal, the 4 videos shown during fear acquisition as well as the 4 videos shown during fear reversal were shown. Hence, fear extinction and fear reversal, in which all CS types were not associated with a

US, only differed in terms of the type of context videos shown, i.e. new context during fear extinction and old, potentially unsafe context during fear renewal.

Cues were presented in a pseudo random order in each experimental phase such that the first and last trials of each participant consisted of unreinforced cues. The assignment of each cue to the 4 possible CS types was counterbalanced across participants as well as the assignment of the type of videos used as context during each experimental phase.

S2: The differential fear learning paradigm involved the presentation of visual stimuli (neutral pictures of an office setting) of which some were paired with electrical stimulation (skin shocks). The stimuli and procedures were modified from Milad et al.[18]. Visual stimuli were presented using the Presentation software package (Neurobehavioral Systems, Berkeley, CA, USA) and MR compatible LCD-goggles (VisuaStim Digital, Resonance Technology, Northridge, CA, USA). Electrical stimulation (1 ms pulses at 50 Hz for a duration of 100 ms) was applied using a constant voltage stimulator (STM2000, BIOPAC Systems, Goleta, CA, USA) along with two electrodes attached to the fingertips of the index and middle finger of the right hand. The intensity of electrical stimulation was adjusted for each participant individually prior to the fear acquisition phase. For this purpose, electrical stimulation was administered at 30 V and raised in increments of 5 V until participants rated the sensation as very unpleasant but not painful.

Each trial started with the presentation of a white fixation cross on a black background for 6.8 - 9.5 seconds. Next, a context image showing an office room with a switched-off desk lamp was presented for 1 second. This was followed by the same image but with the desk lamp either emitting blue (CS+) or yellow light (CS-) for another 6 seconds. In the acquisition phase, the CS+ was paired with electrical stimulation in 62.5 % of trials. The skin shock was administered 5.9 seconds after CS+ onset and co-terminated with CS+ offset. The extinction and renewal phases did not involve any CS+US+ (Fig. 1E, Study 2, first row) presentations but merely CS+US- (Fig. 1E, Study 2, second row) and CS- (Fig. 1E, Study 2, third row) presentations.

During the acquisition phase, CS+ and CS- were presented 16 times each in pseudo-randomized order but distributed equally across both halves of the acquisition phase. The first two trials always involved one CS+ and one CS- presentation and so did the last two trials. The first and last CS+ presentations were always paired with electrical stimulation. The same type of CS was not presented more than twice in consecutive order. The extinction and renewal phases both consisted of 8 CS+US- and 8 CS- presentations.

S3: The study used a differential fear conditioning paradigm based on Ernst et al.[19], using two CSs: black-and-white geometric figures (a square and a diamond) of identical brightness. During fear acquisition training, the CS+ was paired with an aversive US in 62.5% of trials, while the CS- was never followed by the US. The two CS figures were pseudo-randomly counterbalanced across participants. The experiment spanned two days: Day 1 included "habituation" (3 CS+ only trials, 3 CS- only trials), "acquisition training" (10 paired CS+/US trials, 6 CS+ only trials, 16 CS- only trials), and "extinction training" (16 CS+ only trials, 16 CS- only trials). Day 2 featured the recall phase, which was not analysed in the present study. Trial order was pseudorandomized, with specific constraints: the initial two and last trials of the acquisition training were paired CS+/US trials. Additionally, an equal number of events for each type were presented in both the first and second halves of each phase. The sequence of events remained consistent for all participants throughout the habituation, acquisition, and extinction training stages. The electric shock was delivered via a constant current stimulator (DS7A, Digitimer Ltd., London, UK) to the left shin using a concentric bipolar surface electrode with a 6 mm diameter and central platinum pin (WASP electrode, Specialty Developments, Bexley, UK). Electrode position was marked on day 1 for consistency. The US comprised four consecutive 500 µs current pulses with a 33 ms interval, with intensity determined at the start of the experiment. Participants rated sensation intensity until it reached a score of 8 out of 9 (i.e. “unpleasant but not painful”) on a Likert scale. Individual thresholds were increased by 20% to prevent habituation. Each trial involved an 8-second presentation of the CS. In reinforced trials, a 100-millisecond US was presented after 7.9 seconds and ended at the same time as the CS. Intertrial intervals were randomized between 14.3 s and 17.9 s. Each part of the experiment was completed during a different session of fMRI data collection.

S4: The experimental paradigm S4 applied a context-related predictive learning task. Participants learned associations between stimuli/cues (food items) and consequences (occurrence or non-occurrence of a stomach ache) in different contexts (restaurants). After acquisition, the learned associations were extinguished in the extinction phase, and in the final test/recall phase extinction learning and renewal was evaluated. The task contained two conditions defined by the context in which extinction occurred: a) extinction learning in a context different from that present during acquisition and recall (ABA) and b) all learning phases in the same context (AAA). In this way, extinction and recall of stimuli presented in the same and a novel context could be compared and renewal identified.

S5: Conditioned stimuli (CS) consisted of white geometrical shapes (rhomb, parallelogram and square) with similar luminescence on a black background (Fig. 1). To ensure similar luminescence, all white geometrical shapes were designed to match in surface area between stimuli. Each trial had a total duration of 20s and consisted of a black screen presented for 0-2.5s (jittered) at the beginning followed by an 8s presentation of the CS and a jittered 9.5-12s inter-trial interval[20]. The three shapes were randomly assigned to the three CS and balanced between groups. The UCS consisted of a 100ms electrical stimulation applied to the fingertips of the participant’s right index- and middle-finger via two 1cm² electrodes using a constant voltage stimulator (STM200; BIOPAC systems, CA, USA). The stimulation level was adjusted individually to be unpleasant but not painful. The paradigm was presented via MR suitable LCD-goggles (VisuaStim Digital, Resonance Technology Inc., Northridge, CA, USA) and realized in Matlab 2017a (Mathworks Inc., Sherborn, MA, USA).

Fear acquisition training on day one served to establish a CS-UCS association: two of the three CS (CS+G and CS+N) were immediately followed by the UCS in 5/8 trials (62.5% partial reinforcement rate), whereas the CS- was never followed by the UCS (0% reinforcement rate, 8 trials). Importantly, there was no difference in the fear acquisition protocol for CS+G and CS+N (Fig. 1). Prior to fear acquisition training, participants were instructed to pay attention towards possible associations between the presentation of a geometrical shape and the electrical stimulation because they would be asked about them using a questionnaire afterwards. Participants were classified as contingency aware in case they were able to correctly identify the two CS+ (sometimes followed by an electrical stimulation) and the CS- (never followed by an electrical stimulation) after fear acquisition training[20,21]. Before extinction training and retrieval, the participants were informed that the associations learned on day one would not change over the course of the entire experiment to avoid expectancy of contingency reversal. Importantly, participants were not informed about the actual contingencies on any of the three experimental days.

During extinction training on day two, the CS+N and the CS- were presented solely in their original size testing the effects of standard extinction training. The CS+G, however, was additionally to its original size presented in three smaller sizes (75%, 50% and 25% of the original size) to test for the effects of generalised extinction training. The three CS were presented eight times each during extinction training. Each size of the CS+G was presented two times to reach a total number of eight extinction trials; learning effects based on a higher number of presentations (e.g., eight for each size of the CS+G) were avoided using this approach.

During retrieval on day three, all stimuli were presented in their original size for four trials intermixed with four presentations of a formerly unpresented size (175% of the original size) to test for generalisation effects to this new size. Four electrical stimulations, each separated by 5s (after 2s, 7s,

12s and 17s) were applied during the 20s presentation of a grey background for reinstatement after retrieval followed by a reinstatement test (identical to retrieval). After retrieval and before the reinstatement test, a black background was presented for 15s each.

S6: Studies 1 and 2 were two randomized-controlled studies conducted in independent samples of healthy volunteers. Both studies involved two consecutive days, with acquisition training (ACQ) on day 1 and extinction training (EXT) accomplished 24 hours later. Participants were randomized to intravenous injection of lipopolysaccharide (LPS), as an established experimental model of acute systemic inflammation, or to an injection of saline as a placebo. Injections were accomplished two hours before either ACQ (study 1) or EXT (study 2). Paradigms in both studies were identical, with visceral pain induced by pressure-controlled rectal distensions and equally unpleasant, aversive tones implemented as visceral unconditioned stimulus (US+vis) and auditory US (US+aud), respectively, during ACQ. Note that prior to ACQ, US stimulation intensities were individually calibrated and matched for perceived unpleasantness within a predefined perceptual unpleasantness range of 60–80 mm on a 0–100 mm visual analogue scales (based on Koenen et al., 2021). As conditioned stimuli (CS), three distinct visual symbols were paired with either US+vis (CS+US+vis), or US+aud (CS+ US+ aud), or were presented without US (CS-) during ACQ phase. The reinforcement schedule was 75%, with 18 US (9 US+vis, 9 US+aud) and 36 CS presentations (3 out of 12 CS+ presentations were not followed by a US). All CS+ were presented 6–10 s before US (US durations: 14 seconds), with CS and US co-terminating (i.e., differential delay conditioning). During EXT, all CS were presented without US using the same pseudorandomized CS sequence as during ACQ. In all phases, inter-stimulus intervals consisted of a black screen with a fixation cross with a duration of 8 s. Electrodermal activity was continuously recorded using an MRI-compatible system (Biopac Systems, Inc., Goleta, CA, USA; MP160). For analyses reported on herein, only CS+ US+ vis and not CS+ US+ aud were included, and analyses included data from participants treated with LPS or placebo. For methodological details and previously published data from these studies[22,23].

#### Supplemental results

Correlations of learning estimates between FL and PL studies

Splitting these correlations by paradigm (fear learning [FL]: studies S1, S2, S3, S5 and S6; cognitive predictive learning [PL]: study S4) indicated that the correlation between acquisition and extinction was mainly driven by the PL paradigm (FL: $r = .02$, $p_{FDR} > .10$; PL: $r = .35$, $p_{FDR} < .001$; FL vs. PL: $z = 3.03$, $p < .001$). In contrast, acquisition and renewal were negatively correlated in FL ($r = -.38$, $p_{FDR} < .01$), but not in PL ($r = .07$, $p_{FDR} > .10$). Finally, extinction and renewal were not correlated in either FL or PL ($-.13 > r$s $< .09$, $p_{FDR} > .10$).

Our novel procedure for analysing trial-by-trial SCR/behavioural data indicated robust group- and individual-level acquisition, extinction and renewal. Interestingly, a positive association between acquisition and extinction was observed, presumably reflecting the intensity of learning (i.e., participants with higher acquisition levels have more potential for larger decreases in extinction). However, the relationships between learning rates in the different phases differed substantially between FL and PL: For FL, the individual ability to acquire new memories corresponded to a person's renewal. This suggests that participants acquiring high SCRs during initial fear learning also expressed these responses towards the beginning of the renewal phase, but subsequently returned to lower levels; individual abilities of extinction were unrelated, however. In contrast, during PL, individual abilities of acquisition and extinction were related (and may have partly relied on overlapping processes), while renewal was determined by independent factors.

Explained variance for LASSO models

For acquisition, explained variance (deviance ratio) was maximal for FC in PL (mean = 18%), whereas it averaged at 9% and 10 for EC in PL and FL, respectively (also see Fig. S... for a comparison with other resting-state networks). This decrease in explanatory power for FL is probably due to the fact that 1) FL studies were more heterogeneous than PL studies, and 2) the learning variable used in all FL studies was based on SCRs, which is a substantially noisier measure than the behavioural measurements, and also much more diverse among the different FL studies.

For extinction, explained variance for SC was greater for predicting extinction in PL (mean = 22%) than FL (mean = 12%). Explained variance for EC was maximal for PL at 11%, whereas it averaged at 7% in FL. The Poisson regression indicated that for SC, all ROIs were significantly different from zero (zs > 2.10, $p_{FDR}$s $< .05$).

For renewal, explained variance for EC reached 13% in both FLr and PLr.

**References**


1. Esteban, O. *et al.* fMRIPrep: a robust preprocessing pipeline for functional MRI. *Nat. Methods* 16, 111–116 (2019).
2. Markiewicz, C. J. *et al.* fMRIPrep: a robust preprocessing pipeline for functional MRI. Zenodo https://doi.org/10.5281/ZENODO.852659 (2025).
3. Friston, K. J., Kahan, J., Biswal, B. & Razi, A. A DCM for resting state fMRI. *NeuroImage* 94, 396–407 (2014).
4. Beggs, J. M. & Plenz, D. Neuronal Avalanches in Neocortical Circuits. *J. Neurosci.* 23, 11167–11177 (2003).
5. Shin, C.-W. & Kim, S. Self-organized criticality and scale-free properties in emergent functional neural networks. *Phys. Rev. E* 74, 045101 (2006).
6. Stam, C. J. & De Bruin, E. A. Scale-free dynamics of global functional connectivity in the human brain. *Hum. Brain Mapp.* 22, 97–109 (2004).
7. Friston, K., Mattout, J., Trujillo-Barreto, N., Ashburner, J. & Penny, W. Variational free energy and the Laplace approximation. *NeuroImage* 34, 220–234 (2007).
8. Friston, K. J., Harrison, L. & Penny, W. Dynamic causal modelling. *NeuroImage* 19, 1273–1302 (2003).
9. Razi, A. & Friston, K. J. The Connected Brain: Causality, models, and intrinsic dynamics. *IEEE Signal Process. Mag.* 33, 14–35 (2016).
10. Friston, K. J. *et al.* Bayesian model reduction and empirical Bayes for group (DCM) studies. *NeuroImage* 128, 413–431 (2016).
11. Ledoit, O. & Wolf, M. A well-conditioned estimator for large-dimensional covariance matrices. *J. Multivar. Anal.* 88, 365–411 (2004).
12. Giorgino, T. Computing and Visualizing Dynamic Time Warping Alignments in *R* : The dtw Package. *J. Stat. Softw.* 31, (2009).
13. Bach, D. R. A head-to-head comparison of SCRalyze and Ledalab, two model-based methods for skin conductance analysis. *Biol. Psychol.* 103, 63–68 (2014).

14. Staib, M., Castegnetti, G. & Bach, D. R. Optimising a model-based approach to inferring fear learning from skin conductance responses. *J. Neurosci. Methods* 255, 131–138 (2015).

15. Bach, D. R. Psychometrics in experimental psychology: A case for calibration. *Psychon. Bull. Rev.* (2023) doi:10.3758/s13423-023-02421-z.

16. Bach, D. R., Melinščak, F., Fleming, S. M. & Voelkle, M. C. Calibrating the experimental measurement of psychological attributes. *Nat. Hum. Behav.* 4, 1229–1235 (2020).

17. Bach, D. R., Daunizeau, J., Friston, K. J. & Dolan, R. J. Dynamic causal modelling of anticipatory skin conductance responses. *Biol. Psychol.* 85, 163–170 (2010).

18. Bach, D. R., Flandin, G., Friston, K. J. & Dolan, R. J. Modelling event-related skin conductance responses. *Int. J. Psychophysiol.* 75, 349–356 (2010).

19. Bach, D. R., Friston, K. J. & Dolan, R. J. Analytic measures for quantification of arousal from spontaneous skin conductance fluctuations. *Int. J. Psychophysiol.* 76, 52–55 (2010).

20. Milad, M. R. *et al.* Recall of Fear Extinction in Humans Activates the Ventromedial Prefrontal Cortex and Hippocampus in Concert. *Biol. Psychiatry* 62, 446–454 (2007).

21. Ernst, T. M. *et al.* The cerebellum is involved in processing of predictions and prediction errors in a fear conditioning paradigm. *eLife* 8, e46831 (2019).

22. Hagedorn, B., Wolf, O. T. & Merz, C. J. Stimulus-Based Extinction Generalization: Neural Correlates and Modulation by Cortisol. *Int. J. Neuropsychopharmacol.* 24, 354–365 (2021).

23. Tabbert, K. *et al.* Influence of contingency awareness on neural, electrodermal and evaluative responses during fear conditioning. *Soc. Cogn. Affect. Neurosci.* 6, 495–506 (2011).

24. Koenen, Laura. R. *et al.* Associative learning and extinction of conditioned threat predictors across sensory modalities. *Commun. Biol.* 4, 553 (2021).

25. Pawlik, R. J. *et al.* Inflammation shapes neural processing of interoceptive fear predictors during extinction learning in healthy humans. *Brain. Behav. Immun.* 108, 328–339 (2023).

26. Behzadi, Y., Restom, K., Liau, J. & Liu, T. T. A component based noise correction method (CompCor) for BOLD and perfusion based fMRI. *NeuroImage* 37, 90–101 (2007).

27. Greve, D. N. & Fischl, B. Accurate and robust brain image alignment using boundary-based registration. *NeuroImage* 48, 63–72 (2009).

28. Satterthwaite, T. D. *et al.* An improved framework for confound regression and filtering for control of motion artifact in the preprocessing of resting-state functional connectivity data. *NeuroImage* 64, 240–256 (2013).

29. Evans, A. C., Janke, A. L., Collins, D. L. & Baillet, S. Brain templates and atlases. *NeuroImage* 62, 911–922 (2012).

30. Dale, A. M., Fischl, B. & Sereno, M. I. Cortical Surface-Based Analysis. *NeuroImage* 9, 179–194 (1999).

31. Lanczos, C. Evaluation of Noisy Data. *J. Soc. Ind. Appl. Math. Ser. B Numer. Anal.* 1, 76–85 (1964).

32. Pruim, R. H. R. *et al.* ICA-AROMA: A robust ICA-based strategy for removing motion artifacts from fMRI data. *NeuroImage* 112, 267–277 (2015).

33. Jenkinson, M., Bannister, P., Brady, M. & Smith, S. Improved Optimization for the Robust and Accurate Linear Registration and Motion Correction of Brain Images. *NeuroImage* 17, 825–841 (2002).

34. Abraham, A. *et al.* Machine learning for neuroimaging with scikit-learn. *Front. Neuroinformatics* 8, (2014).

35. Power, J. D. *et al.* Methods to detect, characterize, and remove motion artifact in resting state fMRI. *NeuroImage* 84, 320–341 (2014).

36. Klein, A. *et al.* Mindboggling morphometry of human brains. *PLOS Comput. Biol.* 13, e1005350 (2017).

37. Tustison, N. J. *et al.* N4ITK: Improved N3 Bias Correction. *IEEE Trans. Med. Imaging* 29, 1310–1320 (2010).

38. Esteban, O. *et al.* nipy/nipype: 1.9.1. Zenodo https://doi.org/10.5281/ZENODO.596855 (2025).

39. Gorgolewski, K. *et al.* Nipype: A Flexible, Lightweight and Extensible Neuroimaging Data Processing Framework in Python. *Front. Neuroinformatics* 5, (2011).

40. Zhang, Y., Brady, M. & Smith, S. Segmentation of brain MR images through a hidden Markov random field model and the expectation-maximization algorithm. *IEEE Trans. Med. Imaging* 20, 45–57 (2001).

41. Cox, R. W. & Hyde, J. S. Software tools for analysis and visualization of fMRI data. *NMR Biomed.* 10, 171–178 (1997).

42. Avants, B., Epstein, C., Grossman, M. & Gee, J. Symmetric diffeomorphic image registration with cross-correlation: Evaluating automated labeling of elderly and neurodegenerative brain. *Med. Image Anal.* 12, 26–41 (2008).

43. Glasser, M. F. *et al.* The minimal preprocessing pipelines for the Human Connectome Project. *NeuroImage* 80, 105–124 (2013).

44. Fonov, V., Evans, A., McKinstry, R., Almli, C. & Collins, D. Unbiased nonlinear average age-appropriate brain templates from birth to adulthood. *NeuroImage* 47, S102 (2009).

### Supplemental Tables

**Table S1**. Selected studies used in Fig. S36.

| Study | Journal | Amygdala | | Cerebellum | | vmPFC / mPFC | | dorsal Anterior Cingulate Cortex (dACC) | | Hippocampus | |
|---|---|---|---|---|---|---|---|---|---|---|---|
| | | ACQ | EXT | ACQ | EXT | ACQ | EXT | ACQ | EXT | ACQ | EXT |
| Alvarez et al. (2008) | J. Neurosci. | (20,-3,-18) | | | | | | | | (25,-13,-22) | |
| Andreatta et al. (2012) | Learn. Memory | (-32,0,-20) | | | | | | | | | |
| Büchel et al. (1998) | Neuron | (-24,8,-31)<br>(28,1,-32) | | | | | | | | (-24,-42,9)<br>(-27,-22,-6) | |
| Critchley et al. (2002) | Neuron | (17,7,-32)<br>(29,7,-21)<br>(-35,5,30)<br>(-31,-1,-32) | | | | | | | | (27,-7,-35) | |
| Dunasmoor et al. (2011) | Neuroimage | (-30,4,-26)<br>(- 26,4,-18) | | | | | | | | | |
| Eippert et al. (2012) | J. Neurosci. | (29,-7,-17)<br>(-19,-2,-14)<br>(-28,-7,-18)<br>(23,-12,-12)<br>(19,-6,-12)<br>(-22,-4,-14)<br>(-19,-8,-19)<br>(31,-4,-19)<br>(30,-5,-18)<br>(19,0,-18)<br>(21,-8,-18)<br>(18,-6,-14)<br>(-28,-5,-23)<br>(-29,-5,-24)<br>(-31,-2,-21) | | | | | | (-4,10,42)<br>(4,10,34)<br>(-4,10,32)<br>(-4,8,38)<br>(6,16,44)<br>(-6,8,36)<br>(8,12,34)<br>(-4,10,42)<br>(-6,12,30)<br>(-4,12,44) | | | |
| Ernst et al. (2017) | Hum. Brain. Mapp. | | | (38,-51,-28)<br>(-37,-62,-25)<br>(11,-70,-37)<br>(10,-66,-12)<br>(-6,-76,-23)<br>(-10,-40,-28)<br>(11,-40,-28) | (10,-70,-15)<br>(-10,-74,-40)<br>(35,-51,-29)<br>(10,-69,-37)<br>(-33,-55,-32)<br>(-10,-41,-29)<br>(33,-49,-45)<br>(25,-68,-40)<br>(-6,-59,-7) | | | | | | |
| Ewald et al. (2014) | Front. Hum. Neurosci. | | | | | | | (4,22,34) | | | |
| Fani et al. (2014) | Cortex | | | | | | | | | | (-4,36,8 )<br>(-32,-40,-8) |
| Garfinkel et al. (2014) | J. Neurosci. | (-15,-3,-15)<br>(18,0,-15) | (24,-15,-6)<br>(-30,-6,-15) | (-24,-57,-27) | | (0,39,9) | (-3,45,12) | | (15,33,18) | (-12,-33,-6)<br>(9,-33,-6) | (-12,-33,-9)<br>(21,-33,0) |
| Haritha et al. (2012) | PNAS | | | (-2,-59,-32) | | | | | | | |

**Table S1**. (*continuation*)

| Study | Journal | Amygdala | | Cerebellum | | vmPFC / mPFC | | dorsal Anterior Cingulate Cortex (dACC) | | Hippocampus | |
|---|---|---|---|---|---|---|---|---|---|---|---|
| | | ACQ | EXT | ACQ | EXT | ACQ | EXT | ACQ | EXT | ACQ | EXT |
| Hermann et al. (2012) | Plos One | (-27,-4,-26)<br>(24,2,-23)<br>(-30,-4,-20)<br>(30,-1,-26) | | | | (-15,14,-14)<br>(21,11,-17)<br>(-18,11,-14)<br>(18,20,-14) | | (6,8,40) | (-6,11,43) | | |
| Holt et al. (2012) | Arch. Gen. Psychiatry | | | | | | | | | (21,-34,-8)<br>(19,-34,-8) | |
| Icenhour et al. (2015) | Hum. Brain Mapp. | (-12,0,-16) | | | | (32,26,2)<br>(12,26,-20)<br>(6,60,-12) | | | | (-28,-22,-18)<br>(24,-10,-20) | |
| Kalisch et al. (2006) | J. Neurosci. | | | | | | (-2,42,-22) | | | | (-24,-12,-32)<br>(-26,-18,-26) |
| Kattoor et al. (2013) | Plos One | (24,2,-22) | | (16,-68,-12)<br>(32,-66,-18)<br>(-18,-50,-16)<br>(-20,-76,-20)<br>(-38,-50,-26)<br>(6,-72,-10)<br>(-22,-50,-26) | | | (32,6,46)<br>(-10,48,22)<br>(4,54,46)<br>(4,42,24) | | | | |
| Kircher et al. (2013) | Biol. Psychiatry. | (-10,46,-2) | | | | | | | | | |
| Klucken et al. (2012) | Neurosci. | | | | | | | (9,11,40) | | | |
| Kruse et al. (2017) | Soc. Cogn. Affect. Neurosci. | (-24,-2,-12)<br>(20,-2,-12) | (-12,-6,-18)<br>(16,-4,-18) | | | | | (-6,10,42)<br>(4,18,38) | (-10,2,34) | | (-24,38,-4)<br>(22,-38,6) |
| La Bar et al. (1998) | Neuron | | (-17,-2,-17) | | | | | | | | |
| Labrenz et al. (2022) | Behav. Brain Res. | | | (-27,-79,-21)<br>(-32,-62,-30)<br>(-16,-74,-37)<br>(32,-77,-20)<br>(29,-72,-18)<br>(-16,-37,-46)<br>(-7,-56,-43)<br>(-27,-82,-22)<br>(-28,-71,-24)<br>(-10,-38,-22)<br>(-8,-46,-26) | | | | (16,34,22) | | (24,0,-34)<br>(32,-26,-13) | |
| Linnman et al. (2012) | Am. J. Psychiatry | | (-22,-4,-26)<br>(-26,2,-16) | | | | (10,48,-8)<br>(10,42,-18)<br>(-4,48,-18) | | (16,12,30) | | |
| Lissek et al. (2013) | Neuroimage | (-34 ,-4,-28) | | | | | | | | (26,-26,8)<br>(- 18,-24,-12) | |
| Lonsdorf et al. (2014) | Psychopharmacol. | | | | | (0,40,-12) | | | | (29,-16,-25)<br>(36,-32,-14) | |

**Table S1**. (*continuation*)

| Study | Journal | Amygdala | | Cerebellum | | vmPFC / mPFC | | dorsal Anterior Cingulate Cortex (dACC) | | Hippocampus | |
|---|---|---|---|---|---|---|---|---|---|---|---|
| | | ACQ | EXT | ACQ | EXT | ACQ | EXT | ACQ | EXT | ACQ | EXT |
| Lueken et al. (2013) | Psychol. Med. | (-24,-2,-26)<br>(38,4,-26) | | | | | | | (10,54,10) | | |
| Maier et al. (2012) | Plos One | | | | | (-36,48,24)<br>(42,45,18)<br>(-3,48,-18)<br>(-6,44,-20) | | (6,36,33)<br>(3,35,37) | | (-30,-36,-6)<br>(33,-36,-3)<br>(-24,-37,-14)<br>(27,-19,-20) | |
| Marschner et al. (2008) | J. Neurosci. | (22,-6,-26) | | | | | | | | (-34,-24,-14) | |
| Merz et al. (2012) | Soc. Cogn. Affect. Neurosci. | | (33,0,-24)<br>(30,-3,-21)<br>(33,0,-15)<br>(24,3,-24) | | | | (-3,48,-15)<br>(9,57,-3)<br>(-6,48,-15)<br>(9,51,-12)<br>(-3,48,-15)<br>(3,51,-9)<br>(21,18,-12) | | | | |
| Merz et al. (2014) | Soc. Cogn. Affect. Neurosci. | (24,-1,-26)<br>(-21,-13,-14)<br>(21,-13,-14)<br>(-24,-10,-14) | | | | | | | | (33,-22,-11)<br>(-33,-28,-14)<br>(27,-19,-14) | |
| Milad et al. (2007) | Biol. Psychiatry | (26,-5,17) | (-20 ,0 ,-23)<br>(25,-7,-10)<br>(-20,-3,-16) | | (6,-74.5,-43)<br>(12,-75,-47)<br>(11,74,-13)<br>(-11,-74,-47)<br>(-21,81,-48) | (9,19,-16)<br>(9,36,-16) | (8,27,-18)<br>(-10,19,-17) | | | | (-32,-21,-22)<br>(30,-18,-21) |
| Milad et al. (2013) | JAMA Psychiatry | | | | | | (-10,52,-4) | (-10,8,34)<br>(-4,0,54) | (0,14,28)<br>(-4,14,48) | (-34,-28,-14) | |
| Molapour et al. (2015) | Neuroimage | (24,6,-15)<br>(-22,-4,-26) | (- 24,-9,-17) | (-4,-72,-11)<br>(10,-82,-24)<br>(10,-81,-26)<br>(-6,-70,-9)<br>(-14,-60,-29) | | | | | | (-30,-12,-12)<br>(34,-12,-14)<br>(-30,-12,-11)<br>(-24,-42,9)<br>(-27,-22,-6) | (33,-37,4)<br>(-34,-13,-20)<br>(22,-34,6)<br>(18,-13,-18)<br>(-36,-13,-20)<br>(34,-6,-20)<br>(36,-4,-18)<br>(33,-37,4) |
| Olsson et al. (2007) | Soc. Cogn. Affect. Neurosci. | (20,0,-13)<br>(-20,0,-15)<br>(24,1,-16)<br>(-29,-2,-18) | | (5,-50,-26)<br>(40,-54,-31) | | | | (6,37,29)<br>(-1,38,10)<br>(-3,32,27)<br>(-3,13,36)<br>(4,27,35) | | | |
| Pejic et al. (2013) | Soc. Cogn. Affect. Neurosci. | (-12,-6,-18) | (-15,-9,-18)<br>(33,-3,-24) | | | | | (12,15,33) | | (-12,-9,-18) | (-15,-9,-21)<br>(33,-9,-27) |
| Phelps et al. (2004) | Neuron | | (21,-4,-22) | | | | | | (0,19,32) | | (-23,-14,-20)<br>(-23,-14,-20)<br>(22,-12,-17) |

**Table S1**. (*continuation*)

| Study | Journal | Amygdala | | Cerebellum | | vmPFC / mPFC | | dorsal Anterior Cingulate Cortex (dACC) | | Hippocampus | |
|---|---|---|---|---|---|---|---|---|---|---|---|
| | | ACQ | EXT | ACQ | EXT | ACQ | EXT | ACQ | EXT | ACQ | EXT |
| Ploghaus et al. (2000) | PNAS | | | (-25,-65,-35)<br>(23,-62,-34) | | | | | | (-25,-21,-19)<br>(26,-28,-17) | |
| Pohlack et al. (2012) | Biol. Psychol. | (-30,-3,-18)<br>(24,-7,-22) | | (-41,-61,-38) | | | | | | (24,-10,-22) | |
| Ridder et al. (2012) | Psychol. Med. | (-18,0,-12) | | | | | | | | (-27,-12,-21) | |
| Rougemont-Bücking et al. (2011) | CNS Neurosci. Ther. | | | | | | (-6,20,-6) | (-12,26,50) | (10,22,48) | | |
| Schiller et al. (2008) | J. Neurosci. | | | (15,-41,-23) | | | | (2,3,43) | | | |
| Schweckendiek et al. (2010) | Neuroimage | (-24,-12,-12)<br>(-24,-15,-12)<br>(30,0,-15)<br>(-24,-9,-12)<br>(24,-15,-12) | | (-9,-63,-33) | | (3,39,-24)<br>(0,51,-27)<br>(9,42,-12) | | (12,42,21)<br>(12,30,36)<br>(9,6,42) | | | |
| Shvil et al. (2014) | Neurobiol. Learn. Mem. | | | | | (6,38,-16)<br>(-10,38,-18) | | | | | |
| Sripada et al. (2013) | Front. Hum. Neurosci. | (24,3,-24) | (27,3,-27) | | (-9,-87,-33) | | | | | (33,-27,-9)<br>(-21,-30,-3) | (-24,-21,-12)<br>(36,-30,-12) |
| Straube et al. (2014) | Transl. Psychiatry. | | | (12,-68,-16) | | | | (-10,34,26)<br>(-6,42,18) | | | |
| Tabbert et al. (2011) | Soc. Cogn. Affect. Neurosci. | | | | | | | (-3,21,24)<br>(0,6,36)<br>(-3,9,30) | | (-30,-18,-9)<br>(39,-6,-21)<br>(39,-9,-24)<br>(-30,-18,-9)<br>(39,-6,-18) | |
| Utz et al. (2015) | Neurosci. Lett. | | | (35,-44 ,-24) | (7,-47,-11) | | | | | | |
| Wiemer et al. (2015) | Soc. Cogn. Affect. Neurosci. | (-18,-4,-12)<br>(-28,-2,22)<br>(-28,-4,-20) | | (-6,-36,-22)<br>(10,-64,-42)<br>(8,-50,-36)<br>(46,-64,-24)<br>(12,-54,-16) | | (-34,30,36) | | (-2,10,28)<br>(-2,-12,20) | | (-32,-32,-10) | |
| Yágüez et al. (2005) | Gastroenterology | | | (-4,59,-30)<br>(17,-68,-21)<br>(26,72,-14) | | | | | | | |

**Table S2**: Demographics table EHI=Edinburgh Hand Inventory; DASS=Depression, Anxiety and Stress Scale (21 items)

| | Age | Sex | Height (cm) | Weight (kg) | Education (years) | Medication | Hand | EHI | DASS (Depression) | DASS (Anxiety) | DASS (Stress) |
|---|---|---|---|---|---|---|---|---|---|---|---|
| **Mean** | 24.24 | | 1.74 | 70.66 | 15.37 | | | 86.95 | 2.93 | 2.30 | 5.09 |
| **Counts** | | 224 men<br>289 women | | | | 29 | 513 right | | | | |
| **SD** | 4.05 | | 10.96 | 13.36 | 3.20 | | | 18.49 | 3.45 | 2.84 | 4.60 |
| **Range** | 18 - 40 | | 1.49 – 2.00 | 45-120 | 4.5-32 | | | 9.1-100 | 0-24 | 0-12 | 0-26 |

**Table S3.** Event types included in this study. CS=Conditioned stimulus; US=Unconditioned stimulus.

| **Study** | **Acquisition** | **Extinction** | **Study** | **Acquisition** | **Extinction** |
|---|---|---|---|---|---|
| **S1** | **32 x CS+US** | 16 x CS+US | **S2** | **10 x CS+US** | **8 x CS+no US** |
| | **32 x CS+no US** | **48 x CS+no US** | | **6 x CS+no US** | |
| | **64 x CS-** | **48 x CS-** | | **16 x CS-** | **8 x CS-** |
| **S3** | **10 x CS+US** | **16 x CS+no US** | **S4** | **32 x CS+$_{\text{Zum Krug}}$** | 16 x CS+$_{\text{Zum Krug}}$ |
| | **6 x CS+no US** | | | **32 x CS-$_{\text{Zum Krug}}$** | 16 x CS-$_{\text{Zum Krug}}$ |
| | **16 x CS-** | **16 x CS-** | | **32 x CS+$_{\text{Altes Stiftshaus}}$** | 16 x CS+$_{\text{Altes Stiftshaus}}$ |
| | | | | **32 x CS-$_{\text{Altes Stiftshaus}}$** | 16 x CS-$_{\text{Altes Stiftshaus}}$ |
| **S5** | **5 x CS+US$_{gen}$** | 2 x CS+no US$_{gen}$ | **S6** | **9 x CS+US$_{vis}$** | **12 x CS+no US$_{vis}$** |
| | **3 x CS+no US$_{gen}$** | | | **3 x CS+no US$_{vis}$** | |
| | 5 x CS+US$_{ngen}$ | **8 x CS+no US$_{ngen}$** | | 9 x CS+US$_{aud}$ | 12 x CS+no US$_{aud}$ |
| | 3 x CS+no US$_{ngen}$ | | | 3 x CS+no US$_{aud}$ | |
| | **8 x CS-** | **8 x CS-** | | **12 x CS-** | **12 x CS-** |

**Table S4.** Implementation details for each functional connectivity metric used to create the composite metric.

| Metric | Properties | Implementation |
|---|---|---|
| **Pearson's correlation (cor)** | Time-domain; similarity; linear; scale-invariant | $\rho_{cor}(x,y) = \frac{cov(x,y)}{\sqrt{var(x)var(y)}} = \frac{(x-\bar{x})(y-\bar{y})^T}{\left(\sqrt{(x-\bar{x})(x-\bar{x})^T}\right)\{\sqrt{((y-\bar{y})(y-\bar{y})^T)}\}}$ |
| **Cross-correlation (xcor)** | Time-domain; similarity; linear; scale-invariant | $\rho_{xcor}(x,y) = \rho_{xy}(m) = \begin{Bmatrix} \sum_{i=0}^{t-m-1} x_{i+m} y_i^* \; if\ m \geq 0 \\ \rho_{yx}(-m) \quad if\ m < 0 \end{Bmatrix}$ |
| **Coherence (cohe)** | Frequency-domain; similarity; linear; scale-invariant | $\rho_{cohe}(x,y) = \frac{\lvert P_{xy}(f)\rvert^2}{P_{xx}(f)P_{yy}(f)}$ |
| **Wavelet-coherence (wcohe)** | Time-frequency domain; similarity; linear; scale-invariant | $\rho_{wcohe}(x,y) = \frac{\lvert S(C_x(a,b)C_y(a,b))\rvert^2}{S(\lvert C_x(a,b)\rvert^2)S(\lvert C_x(a,b)\rvert^2)}$ |
| **Mutual information (MI)** | Time-domain; similarity; non-linear; scale-variant | $\rho_{MI}(x,y) = I(x;y) = \sum_{y \in Y=0} \sum_{x \in X} p(x,y)\, log \frac{p(x,y)}{p(x)p(y)}$ |
| **Euclidean distance (ED)** | Time-domain; dissimilarity; linear; scale-variant | $\rho_{ED}(x,y) = \lvert\lvert x-y \rvert\rvert_2 = \sqrt{(x-y)(x-y)^T}$ |
| **Manhattan (citiblock) distance (MD)** | Time-domain; dissimilarity; linear; scale-variant | $\rho_{MD}(x,y) = \lvert\lvert x-y \rvert\rvert_1 = \sum_{i=1}^{t} \lvert x_i - y_i \rvert$ |
| **Dynamic time warping (DTW)** | Time-domain; dissimilarity; non-linear; scale-variant | $\rho_{DTW}(x,y) = min_{\pi \epsilon A(x,y)} \sum_{(i,j)\epsilon\pi} d(x_i, y_j)^2$ |
| **Wasserstein (Earth mover's) distance (WD)** | Time-domain; dissimilarity; non-linear; scale-variant | $\rho_{WD}(x,y) = \frac{\sum_{i=1}^{m}\sum_{j=1}^{n} f_{ij} d_{ij}}{\sum_{i=1}^{m}\sum_{j=1}^{n} f_{ij}}$ |

**Table S5**. Variance estimates of the random effects “study” and “experiment” after standardising the learning variable were zero, indicating that that the model with random effects over and above “participant” was degenerate, and, thus, their inclusion is not justifiable.

| **Group** | **Variance** |
|---|---|
| Experiment:(Study:Participant) | 0.0 |
| Study:Participant | 0.0 |
| Participant | 0.0 |
| Residual | 0.99 |

**Table S6**. Model comparisons using different random effects. Adding more random effects to the model beyond "Participant" does not improve model fit.

| Group | Variance | $\chi^2$ | D.O.F | Pr (> $\chi^2$) |
|---|---|---|---|---|
| Participant | 3092.9 | | | |
| Participant + Study | 3094.9 | 0 | 1 | 1 |
| Participant + Study + Experiment | 3096.9 | 0 | 1 | 1 |
| Participant + Study + Experiment + Site | 3096.9 | 0 | 1 | 1 |

**Table S7**. No systematic "site" effects present in the data. Comparisons between the original model before ComBat and the model after ComBat correction led to negligible differences between the two models.

| Modality | Phase | Original model | ComBat model | ΔAIC |
|---|---|---|---|---|
| FC | Acquisition | 1140.15 | 1140.10 | < 0.1 |
| FC | Extinction | 1074.04 | 1074.00 | < 0.1 |
| SC | Acquisition | 1041.45 | 1041.44 | < 0.1 |
| SC | Extinction | 983.77 | 983.76 | < 0.1 |
| EC | Acquisition | 1084.58 | 1084.58 | < 0.1 |
| EC | Extinction | 1160.74 | 1160.65 | < 0.1 |

**Table S8**. Grouping codes used in lasso models and their corresponding description. Bold letters in the "Description" column indicate how the acronyms in the "Grouping code" column were constructed.

| Grouping code | Groups | # of experiments | Description |
|---|---|---|---|
| PL | S4 | 4 | **P**redictive **L**earning |
| PLr | S4 | 4 | **P**redictive **L**earning (**r**enewal only) |
| FLr | S2 | 2 | **F**ear **L**earning (**r**enewal only) |
| FLc | S2 S3 | 3 | **F**ear **L**earning using a **c**lassical paradigm and same PsPM analysis method |
| FLs | S2 S3 S5 | 5 | **F**ear **L**earning using a **s**imilar paradigm and same PsPM analysis method |
| Flel | S1 S2 S3 S5 | 6 | **F**ear **L**earning studies that used **el**ectric shocks as US |
| FLst | S2 S3 S5 S6 | 7 | **F**ear **L**earning studies that employed a **st**andard extinction learning experimental procedure |
| FL | S1 S2 S3 S5 S6 | 8 | All **F**ear **L**earning studies |
| All | S1 S2 S3 S4 S5 S6 | 12 | **All** studies |

**Table S9.** Pairwise comparisons among connections regarding functional connectivity strength. ACC=Dorsal anterior cingulate cortex; AMY=Amygdala; CEB=Cerebellar nuclei; HIP=Hippocampus; PFC=Ventro-medial prefrontal cortex. Significance levels: * $p < .05$, ** $p < .01$, *** $p < .001$

| Connection 1 | Connection 2 | Estimate | T ratio | Significance |
|---|---|---|---|---|
| rACC-rPFC | rHIP-rACC | 0,05 | 2,10 | * |
| rACC-rPFC | rHIP-rPFC | -0,55 | -23,58 | *** |
| rAMY-rACC | rAMY-rHIP | -1,25 | -53,95 | *** |
| rAMY-rACC | rAMY-rPFC | -0,22 | -9,34 | *** |
| rAMY-rACC | rHIP-rACC | 0,00 | -0,09 | n.s. |
| rAMY-rACC | rHIP-rPFC | -0,60 | -25,77 | *** |
| rAMY-rHIP | rAMY-rPFC | 1,04 | 44,61 | *** |
| rAMY-rHIP | rHIP-rACC | 1,25 | 53,86 | *** |
| rAMY-rHIP | rHIP-rPFC | 0,66 | 28,18 | *** |
| rAMY-rPFC | rHIP-rACC | 0,22 | 9,25 | *** |
| rAMY-rPFC | rHIP-rPFC | -0,38 | -16,42 | *** |
| rHIP-rACC | rHIP-rPFC | -0,60 | -25,68 | *** |
| lAMY-rCEB | lHIP-lACC | 0,13 | 5,43 | *** |
| lAMY-rCEB | lHIP-lPFC | -0,59 | -25,29 | *** |
| lAMY-rCEB | rCEB-lACC | 0,06 | 2,48 | * |
| lAMY-rCEB | rCEB-lHIP | 0,06 | 2,54 | * |
| lAMY-rCEB | rCEB-lPFC | 0,11 | 4,79 | *** |
| rCEB-lACC | rCEB-lHIP | 0,00 | 0,06 | n.s. |
| lCEB-rHIP | rAMY-rACC | 0,02 | 0,78 | n.s. |
| lCEB-rHIP | rAMY-rHIP | -1,24 | -53,17 | *** |
| lCEB-rHIP | rAMY-rPFC | -0,20 | -8,57 | *** |
| lCEB-rHIP | rHIP-rACC | 0,02 | 0,69 | n.s. |
| lCEB-rHIP | rHIP-rPFC | -0,58 | -24,99 | *** |
| lCEB-rPFC | rACC-rPFC | -0,01 | -0,29 | n.s. |
| lCEB-rPFC | rAMY-lCEB | 0,06 | 2,46 | * |
| lCEB-rPFC | rAMY-rACC | 0,04 | 1,90 | n.s. |
| lCEB-rPFC | rAMY-rHIP | -1,21 | -52,05 | *** |
| lCEB-rPFC | rAMY-rPFC | -0,17 | -7,45 | *** |
| lCEB-rPFC | rHIP-rACC | 0,04 | 1,81 | n.s. |
| lCEB-rPFC | rHIP-rPFC | -0,56 | -23,87 | *** |
| rAMY-lCEB | rAMY-rACC | -0,01 | -0,56 | n.s. |
| rAMY-lCEB | rAMY-rHIP | -1,27 | -54,51 | *** |
| rAMY-lCEB | rAMY-rPFC | -0,23 | -9,90 | *** |
| rAMY-lCEB | rHIP-rACC | -0,02 | -0,65 | n.s. |
| rAMY-lCEB | rHIP-rPFC | -0,61 | -26,33 | *** |

**Table S10.** Pairwise comparisons among connections regarding structural connectivity strength. ACC=Dorsal anterior cingulate cortex; AMY=Amygdala; CEB=Cerebellar nuclei; HIP=Hippocampus; PFC=Ventro-medial prefrontal cortex. Significance levels: * $p < .05$, ** $p < .01$, *** $p < .001$

| Connection 1 | Connection 2 | Estimate | T ratio | Significance |
|---|---|---|---|---|
| lACC-lPFC | lAMY-lACC | 5,44 | 86,36 | *** |
| lACC-lPFC | lAMY-lHIP | -3,81 | -60,52 | *** |
| lACC-lPFC | lAMY-lPFC | 1,64 | 26,08 | *** |
| lACC-lPFC | lHIP-lACC | 4,00 | 63,50 | *** |
| lACC-lPFC | lHIP-lPFC | 2,41 | 38,33 | *** |
| lAMY-lACC | lAMY-lHIP | -9,25 | -146,88 | *** |
| lAMY-lACC | lAMY-lPFC | -3,79 | -60,28 | *** |
| lAMY-lACC | lHIP-lACC | -1,44 | -22,86 | *** |
| lAMY-lACC | lHIP-lPFC | -3,02 | -48,03 | *** |
| lAMY-lHIP | lAMY-lPFC | 5,45 | 86,60 | *** |
| lAMY-lHIP | lHIP-lACC | 7,81 | 124,02 | *** |
| lAMY-lHIP | lHIP-lPFC | 6,22 | 98,85 | *** |
| lAMY-lPFC | lHIP-lACC | 2,36 | 37,42 | *** |
| lAMY-lPFC | lHIP-lPFC | 0,77 | 12,25 | *** |
| lHIP-lACC | lHIP-lPFC | -1,58 | -25,17 | *** |
| lAMY-rCEB | lHIP-lACC | -1,21 | -19,24 | *** |
| lAMY-rCEB | lHIP-lPFC | -2,80 | -44,41 | *** |
| lAMY-rCEB | rCEB-lACC | 2,24 | 35,56 | *** |
| lAMY-rCEB | rCEB-lHIP | 0,03 | 0,46 | n.s |
| lAMY-rCEB | rCEB-lPFC | 1,77 | 28,04 | *** |
| rCEB-lACC | rCEB-lHIP | -2,21 | -35,10 | *** |
| rCEB-lACC | rCEB-lPFC | -0,47 | -7,52 | *** |
| rCEB-lHIP | rCEB-lPFC | 1,74 | 27,58 | *** |
| rACC-rPFC | rAMY-rACC | 5,24 | 83,19 | *** |
| rACC-rPFC | rAMY-rHIP | -3,49 | -55,48 | *** |
| rACC-rPFC | rAMY-rPFC | 2,95 | 46,88 | *** |
| rACC-rPFC | rHIP-rACC | 3,73 | 59,29 | *** |
| rACC-rPFC | rHIP-rPFC | 4,15 | 65,86 | *** |
| rAMY-rACC | rAMY-rHIP | -8,73 | -138,68 | *** |
| rAMY-rACC | rAMY-rPFC | -2,29 | -36,31 | *** |
| rAMY-rACC | rHIP-rACC | -1,50 | -23,90 | *** |
| rAMY-rACC | rHIP-rPFC | -1,09 | -17,33 | *** |
| rAMY-rHIP | rAMY-rPFC | 6,44 | 102,36 | *** |

**Table S10.** (*continuation*)

| Connection 1 | Connection 2 | Estimate | T ratio | Significance |
|---|---|---|---|---|
| rAMY-rHIP | rHIP-rACC | 7,23 | 114,77 | *** |
| rAMY-rHIP | rHIP-rPFC | 7,64 | 121,34 | *** |
| rAMY-rPFC | rHIP-rACC | 0,78 | 12,41 | *** |
| rAMY-rPFC | rHIP-rPFC | 1,19 | 18,98 | *** |
| rHIP-rACC | rHIP-rPFC | 0,41 | 6,57 | *** |
| lCEB-rACC | lCEB-rHIP | -2,16 | -34,25 | *** |
| lCEB-rACC | lCEB-rPFC | 1,11 | 17,66 | *** |
| lCEB-rACC | rACC-rPFC | -7,54 | -119,83 | *** |
| lCEB-rACC | rAMY-lCEB | -2,43 | -38,54 | *** |
| lCEB-rACC | rAMY-rACC | -2,31 | -36,63 | *** |
| lCEB-rACC | rAMY-rHIP | -11,04 | -175,31 | *** |
| lCEB-rACC | rAMY-rPFC | -4,59 | -72,94 | *** |
| lCEB-rACC | rHIP-rACC | -3,81 | -60,54 | *** |
| lCEB-rACC | rHIP-rPFC | -3,40 | -53,97 | *** |
| lCEB-rHIP | lCEB-rPFC | 3,27 | 51,92 | *** |
| lCEB-rHIP | rACC-rPFC | -5,39 | -85,57 | *** |
| lCEB-rHIP | rAMY-lCEB | -0,27 | -4,29 | ** |
| lCEB-rHIP | rAMY-rACC | -0.15 | -2,38 | n.s. |
| lCEB-rHIP | rAMY-rHIP | -8,88 | -141,05 | *** |
| lCEB-rHIP | rAMY-rPFC | -2,44 | -38,69 | *** |
| lCEB-rHIP | rHIP-rACC | -1,65 | -26,28 | *** |
| lCEB-rHIP | rHIP-rPFC | -1,24 | -19,71 | *** |
| lCEB-rPFC | rACC-rPFC | -8,66 | -137,49 | *** |
| lCEB-rPFC | rAMY-lCEB | -3,54 | -56,21 | *** |
| lCEB-rPFC | rAMY-rACC | -3,42 | -54,29 | *** |
| lCEB-rPFC | rAMY-rHIP | -12,15 | -192,97 | *** |
| lCEB-rPFC | rAMY-rPFC | -5,70 | -90,61 | *** |
| lCEB-rPFC | rHIP-rACC | -4,92 | -78,20 | *** |
| lCEB-rPFC | rHIP-rPFC | -4,51 | -71,63 | *** |
| rAMY-lCEB | rAMY-rACC | 0,12 | 1,91 | *** |
| rAMY-lCEB | rAMY-rHIP | -8,61 | -136,76 | *** |
| rAMY-lCEB | rAMY-rPFC | -2,17 | -34,40 | *** |
| rAMY-lCEB | rHIP-rACC | -1,38 | -21,99 | *** |
| rAMY-lCEB | rHIP-rPFC | -0,97 | -15,42 | *** |

**Table S11.** Model performance for each experimental phase.

| Phase | Difference | Statistic | Low C.I. | High C.I. | P-value |
|---|---|---|---|---|---|
| Acquisition | .05 | 5.80 | -.06 | -.03 | < .001 |
| Extinction | .05 | 5.80 | -.06 | -.03 | < .001 |
| Renewal | .05 | 7.73 | -.07 | -.04 | < .001 |

#### Supplemental Figures

**Figure S1**. **(A)** The fifty-six regions of interest (ROIs) selected for the analysis of the travelling heads. **(B)** Establishment of test-retest reliability across scanning sites. Two individuals (travel heads) were scanned on three different scanners over the course of three years (3 sessions). Functional connectivity (FC) for each pair of ROIs (top panel), and fractional anisotropy (FA) within each ROI (bottom panel) were computed for every session, site and travel head. Left: Test-retest reliability was very high across all sessions, but also for each site and travel head. Middle: Correlations within and between sites were highly significant for both FC and FA. Right: FC and FA estimates of each ROI (coloured lines) remained largely stable across the different sessions.

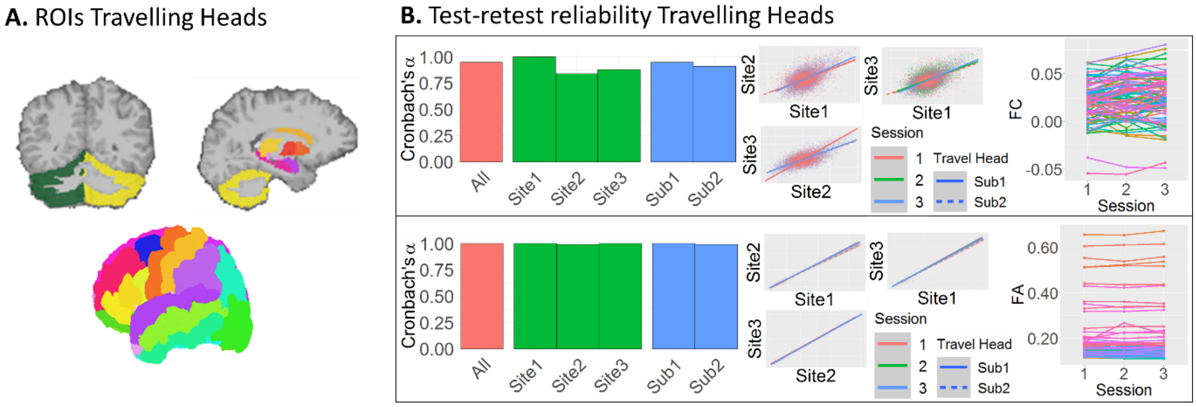

**Figure S2**. Same as Figure S1B, but only for the ROIs used in the main study (amygdala, hippocampus, dorsal anterior cingulate cortex, ventro-medial prefrontal cortex, cerebellar nuclei).

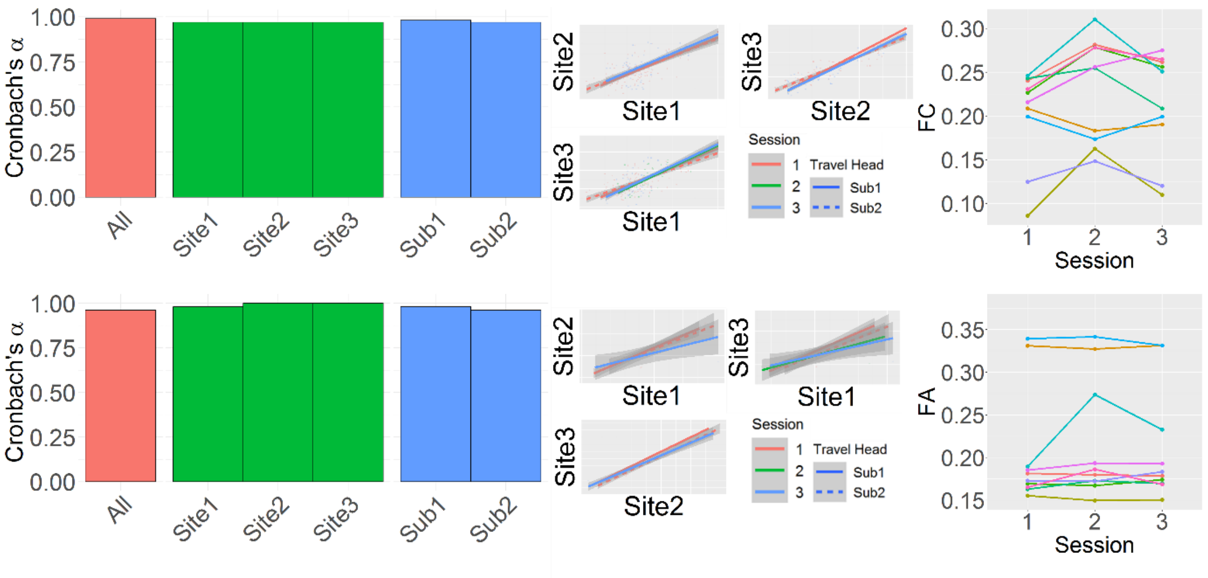

**Figure S3**. Experimental paradigms of each group. S1: During acquisition, a context image was followed by either a CS+ item (partial reinforcement with US skin shock administered after 2.5 seconds) or a CS- item. During extinction, half of all items retained their contingencies, for the other half contingencies were reversed. S2: During acquisition, a context image was followed by either a CS+ item (partial reinforcement with US skin shock after 5.9 seconds) or a CS- item. During extinction, CS- and CS+US- items were shown. S3: Similar to S2 but without a context image and with shapes instead of images as CSs. S4: Subjects indicated whether a food item predicted stomach ache during acquisition. During extinction, outcomes reversed for half of the food items. S5: Similar to S3 with the exception that CS+US- items were presented in three different sizes during extinction. S6: During acquisition, distinct visual CS were paired with either an aversive tone or a moderately painful rectal distension serving as US. During extinction, CS+US- and CS- were shown.

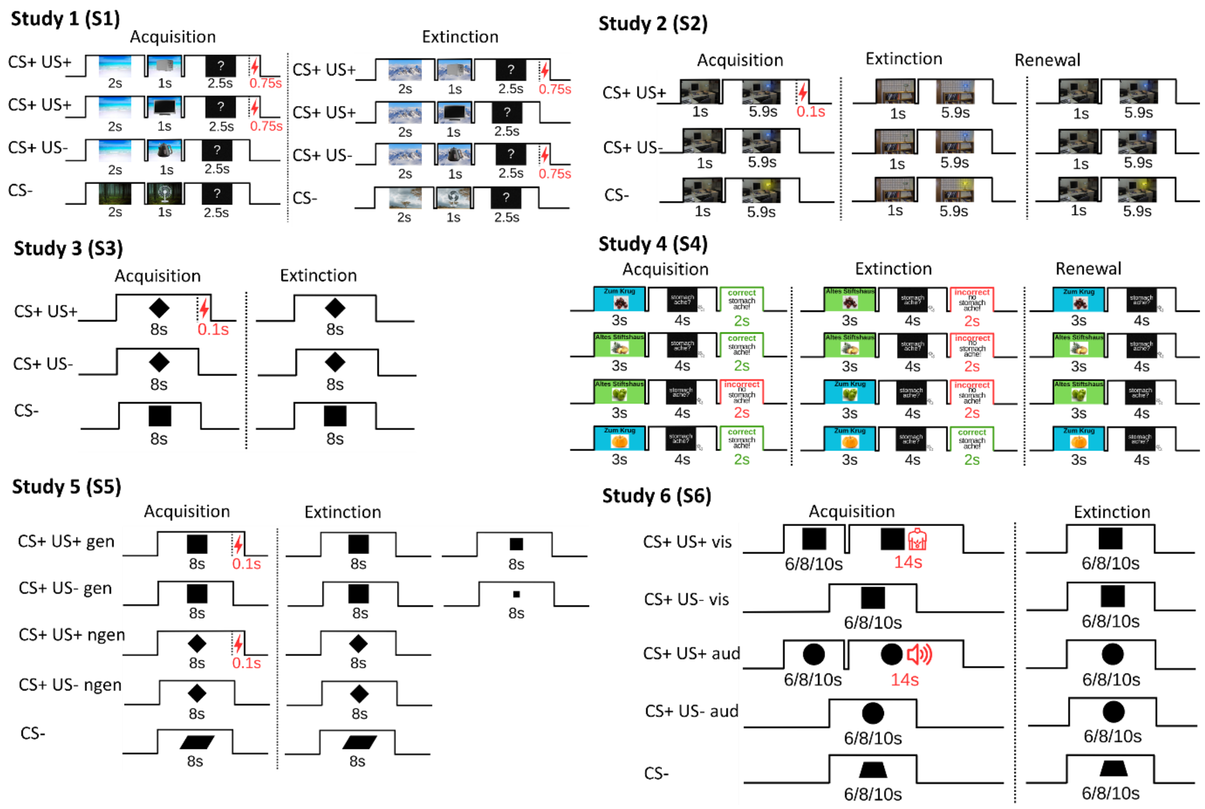

**Figure S4**. For probabilistic tractography, surfaces of the dorsal anterior cingulate and ventro-medial prefrontal cortices were used instead of their volumetric counterparts.

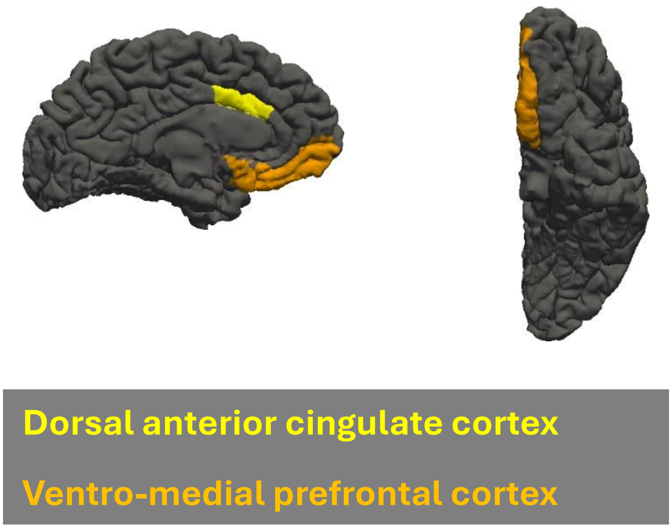

**Figure S5**. Schematic illustration of the pipelines used in the present study.

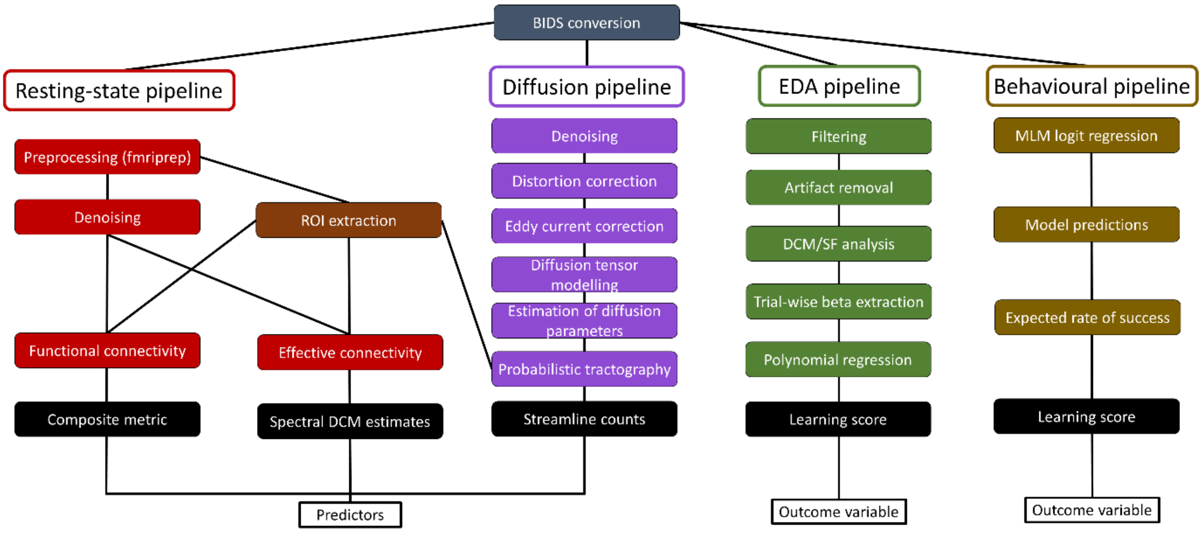

**Figure S6**. Reconstructed tracts for the connections between the hippocampus (HIP) and amygdala (AMY), dorsal anterior cingulate cortex (ACC), cerebellar nuclei (CEB) and ventro-medial prefrontal cortex (PFC).

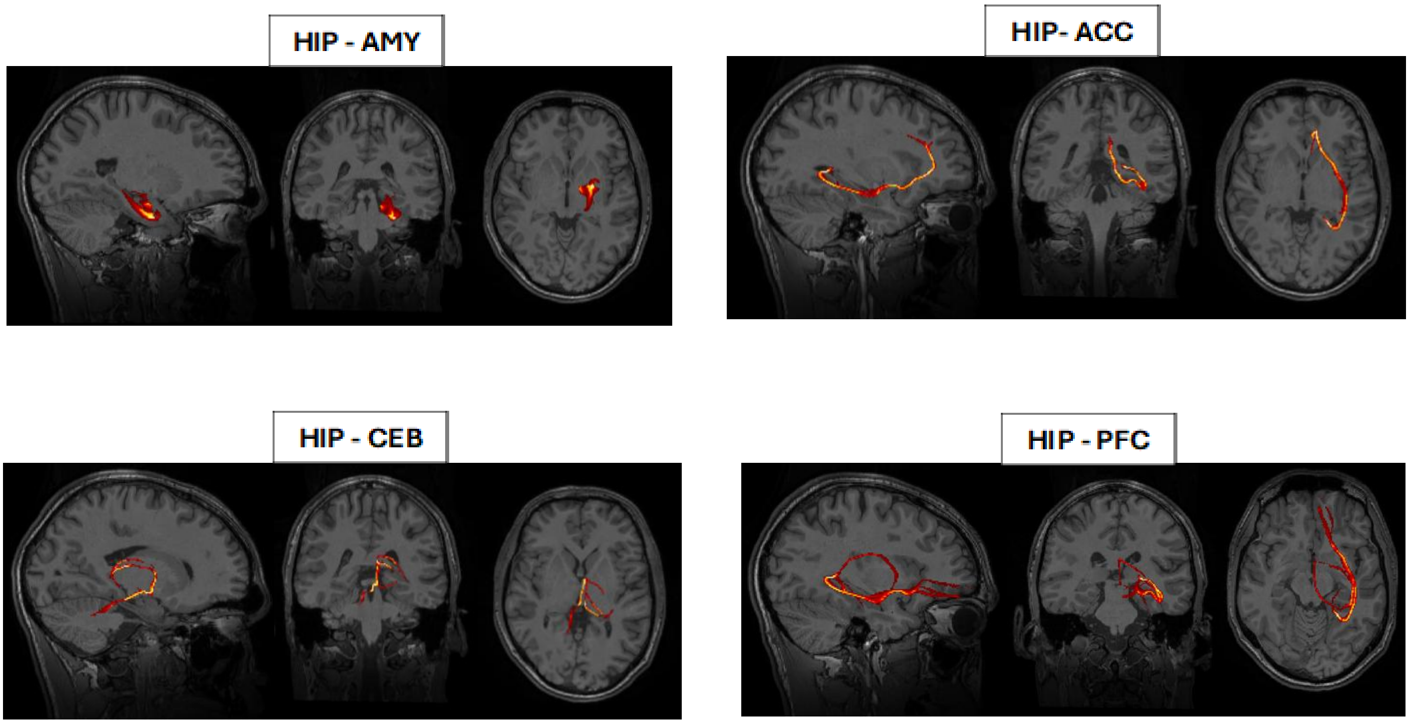

**Figure S7**. Correlations among functional connectivity metrics. All metrics were highly correlated with the Composite score (Comp). In addition, most metrics were correlated among themselves with the exception of Wasserstein (Cityblock) distance for some studies. Cor=Pearson correlation; Xcor = cross-correlation; ED=Euclidean distance; MD=Manhattan distance; WD=Wasserstein distance; Dtw=Dynamic time warping; MI=Mutual information; Cohe=Coherence; Wcohe=Wavelet coherence; Comp=Composite.

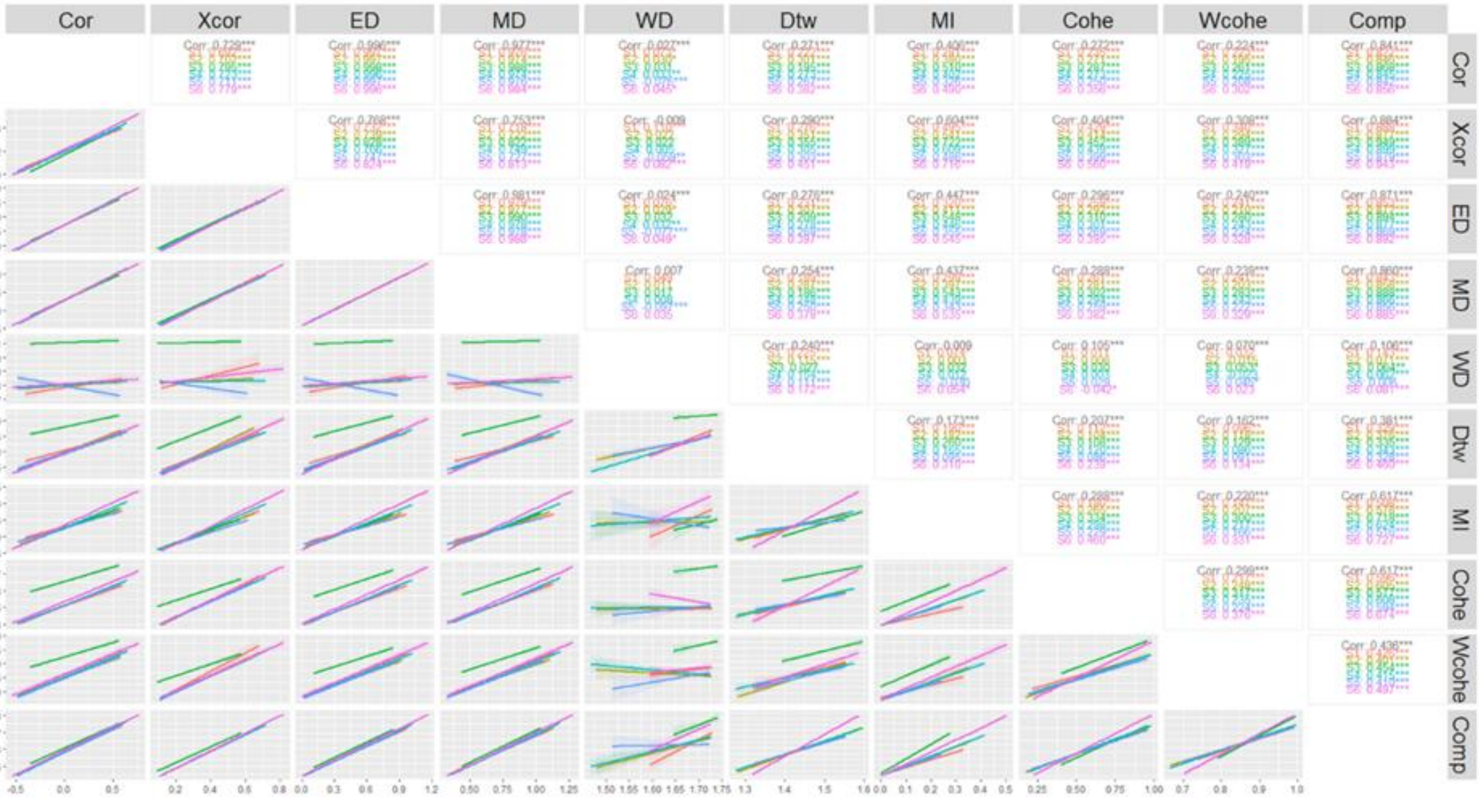

**Figure S8**: Functional connectivity within the extinction learning network regions for each of the nine metrics separately. CEB = cerebellar nuclei, HIP = hippocampus, AMY = amygdala, ACC = dorsal anterior cingulate cortex, PFC = ventro-medial prefrontal cortex.

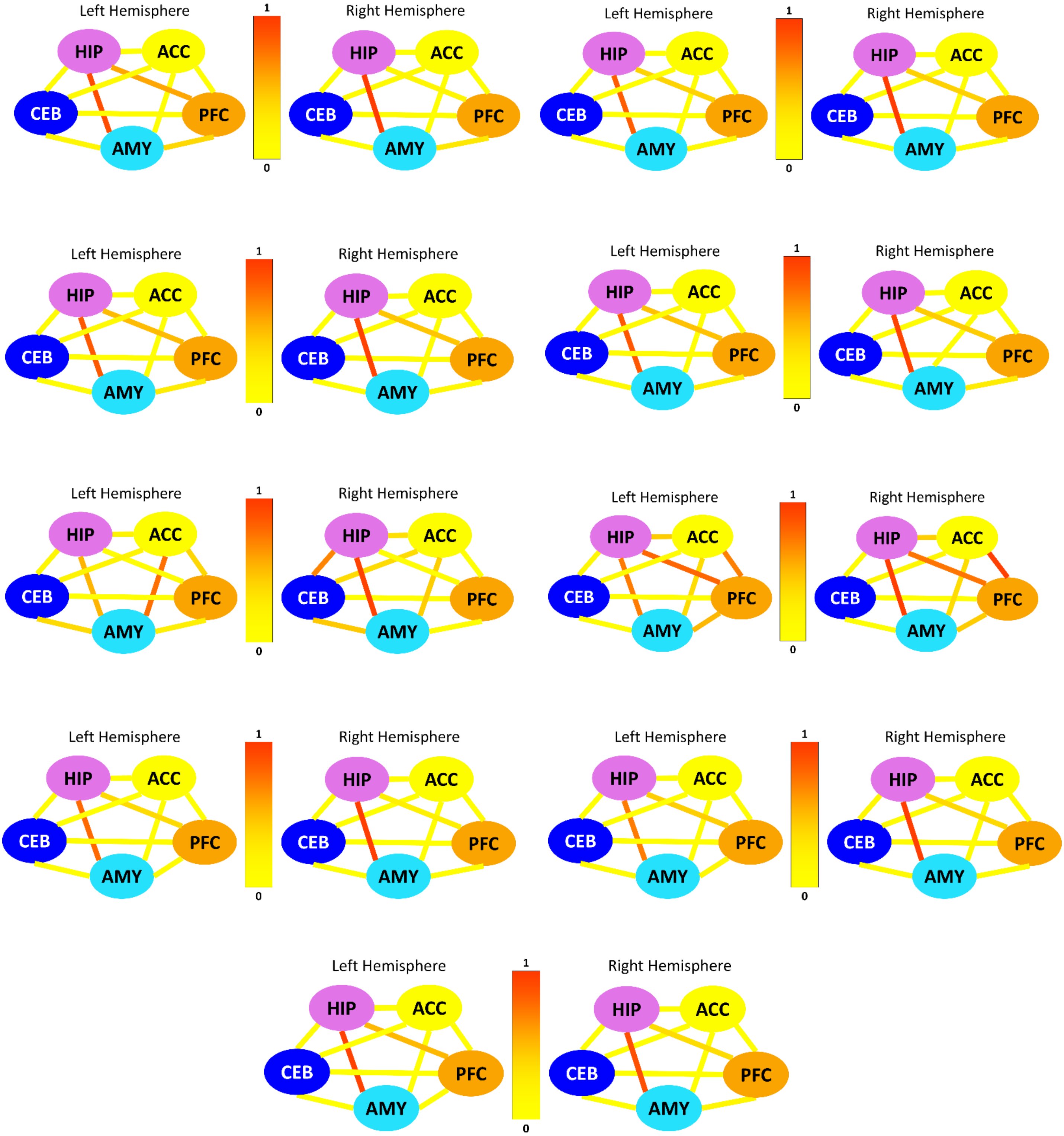

**Figure S9**. Top: Average correlation across ROIs among the different functional connectivity metrics. Bottom: Normalised correlation values for each individual connectivity pair and functional connectivity metric. Corr=Pearson correlation; Xcorr=Cross-correlation; ED=Euclidean distance; MD=Manhattan distance; WD=Wasserstein distance; DTW=Dynamic time warping; MI=Mutual information; Cohe=Coherence; wCohe=Wavelet coherence; Comp=Composite.

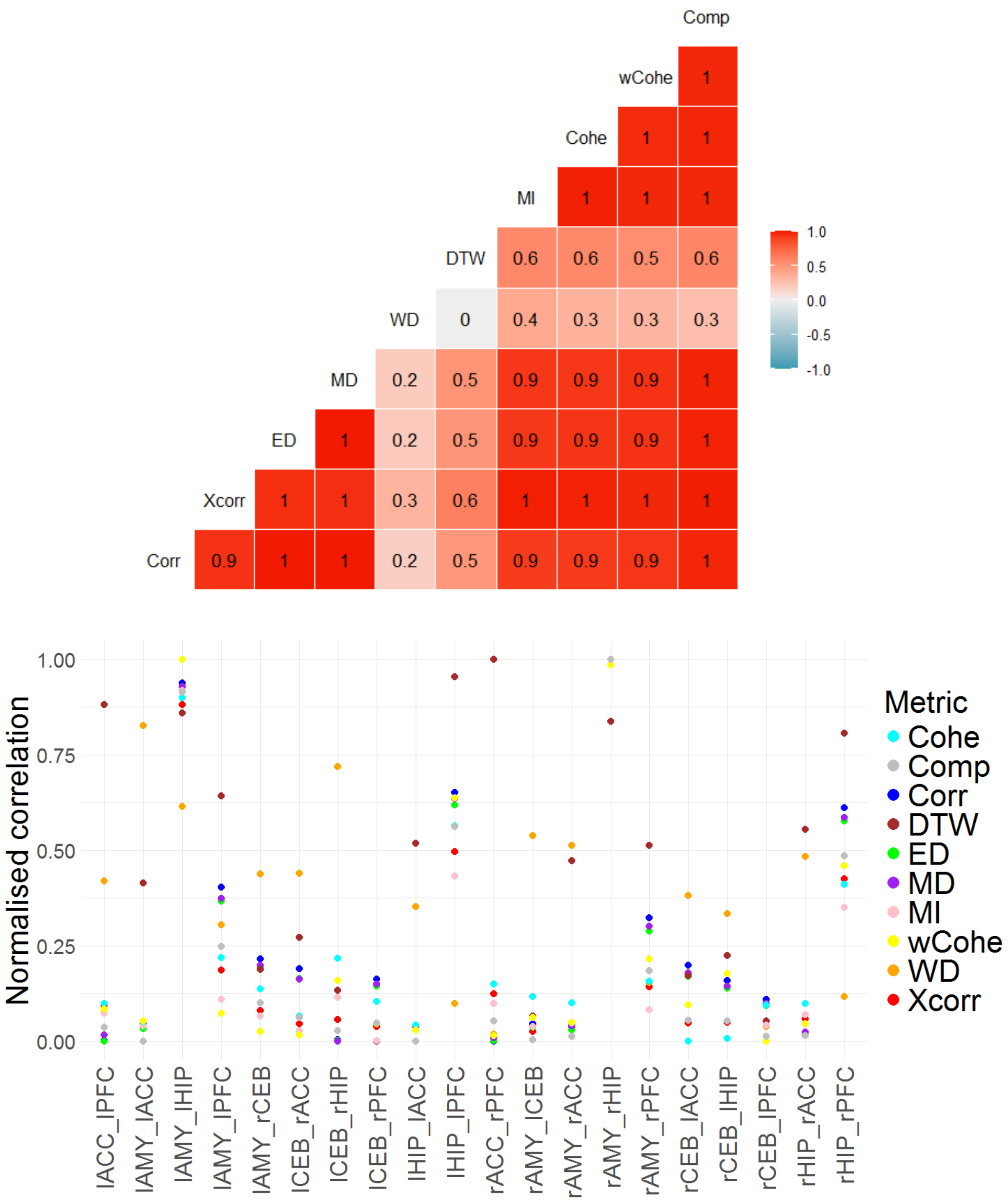

**Figure S10.** Correlation between spectral DCM (spDCM) estimates and coherence (left) and cross-correlation (right). Stars show the significant pairs after correcting for multiple comparisons.

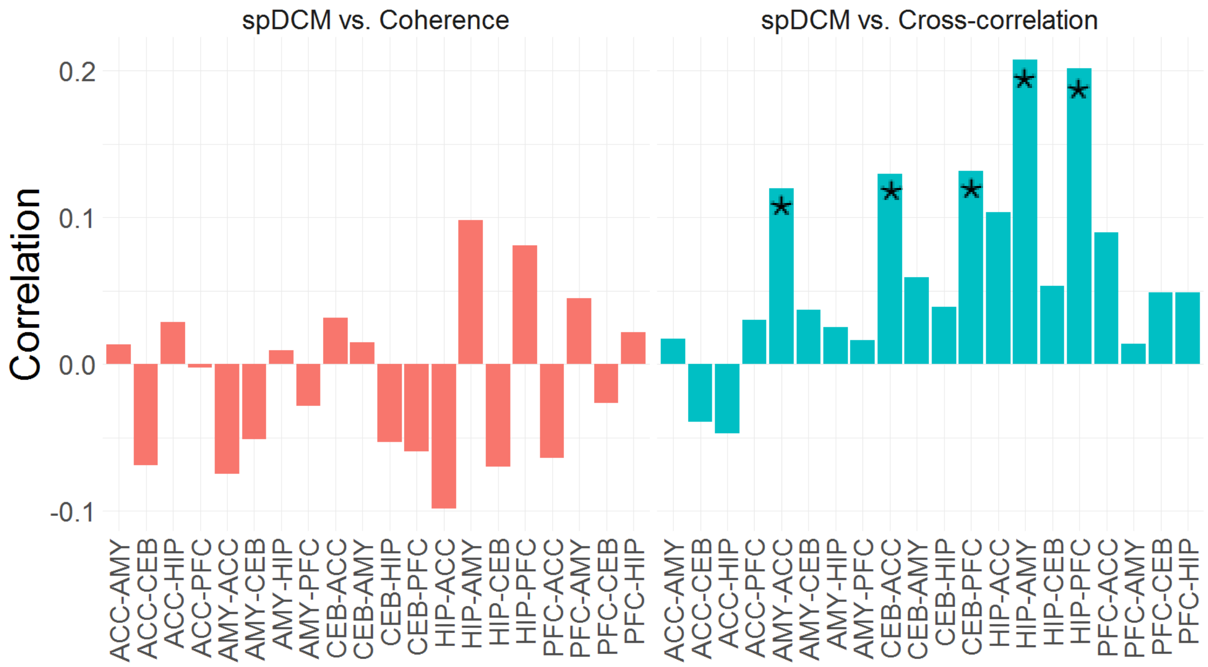

**Figure S11**. Significant connections showing greater connectivity for FC > SC (red) or SC > FC (blue) within the fear and extinction network regions. CEB = cerebellar nuclei, HIP = hippocampus, AMY = amygdala, ACC = dorsal anterior cingulate cortex, PFC = ventro-medial prefrontal cortex. FC = Functional connectivity; SC = Structural connectivity; EC = Effective connectivity.

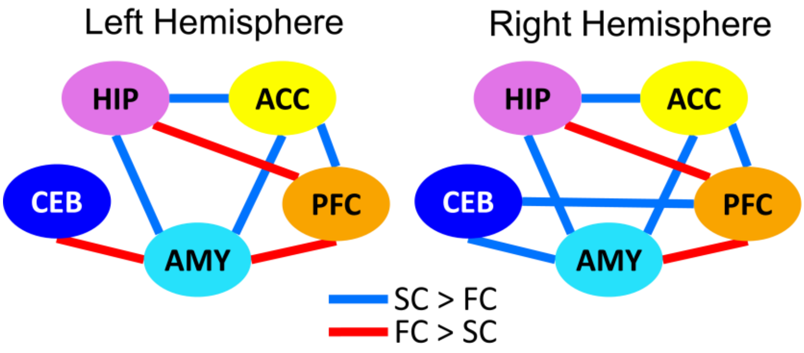

**Figure S12**. Correlations among the different connectivity modalities. Only those connections that survived correction for multiple comparisons are shown. CEB = cerebellar nuclei, HIP = hippocampus, AMY = amygdala, ACC = dorsal anterior cingulate cortex, PFC = ventro-medial prefrontal cortex. FC = Functional connectivity; SC = Structural connectivity; EC = Effective connectivity.

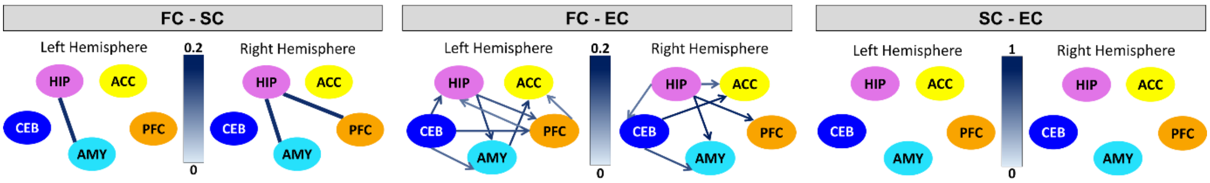

**Figure S13**. Examples of excluded EDA datasets

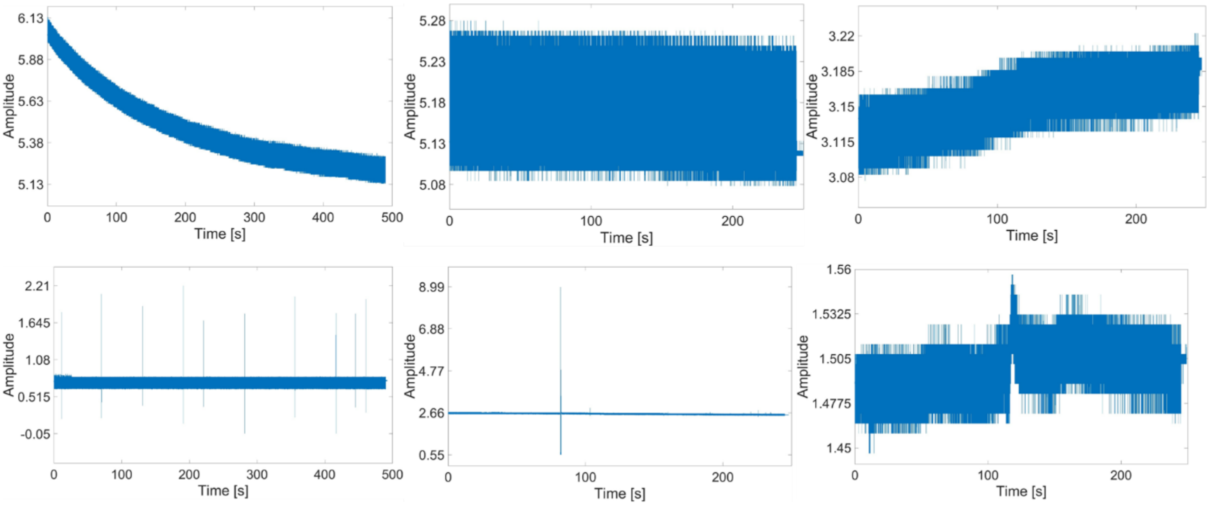

**Figure S14**. Mean sudomotor response latency of SCR values during the acquisition phase was below the expected value (the estimated onset of the Gaussian bump plus at least 2 standard deviations; red lines) had the US contaminated the CS responses. This check was unnecessary for extinction and renewal phases, since no shock was administered. Also, even though we also analysed SCR for S6, the approach we used for that study ensured that US contamination was not possible (see main text).

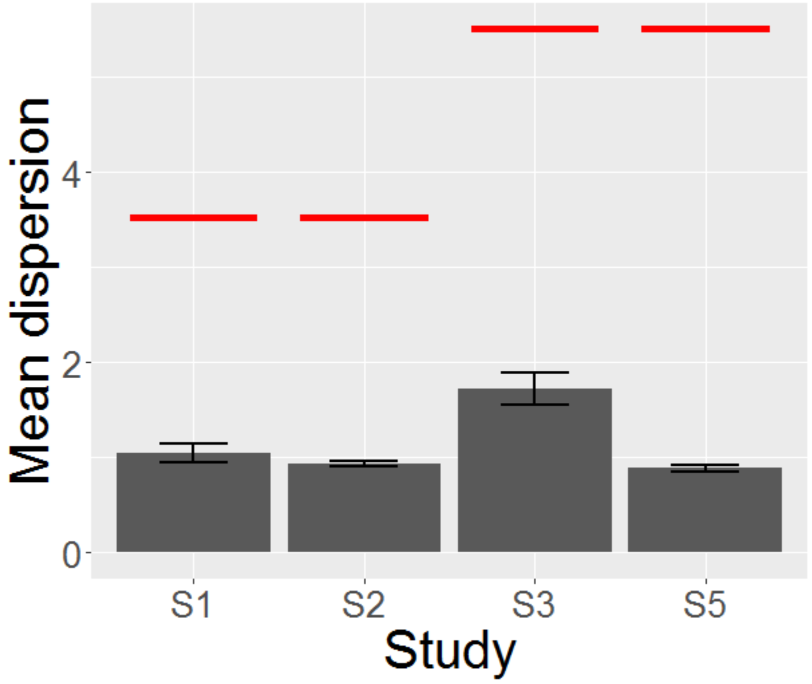

**Figure S15**. Expected learning success for each participant (dots) based on the model shown in Figure 2D in the main paper. Dashed line indicates chance level. Error bars represent standard error of the mean.

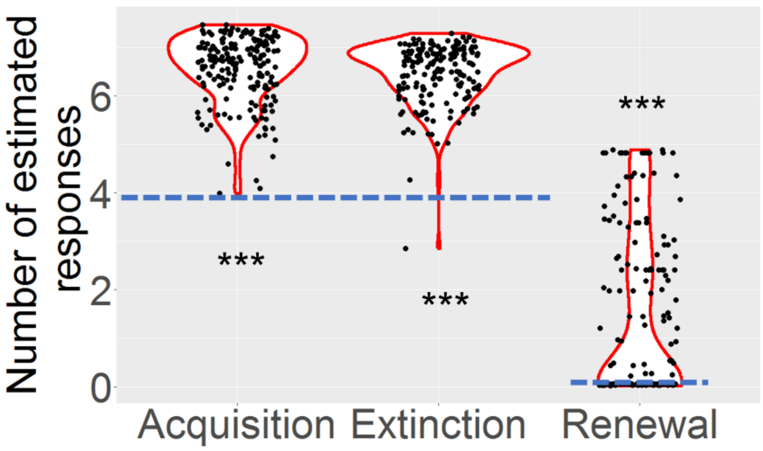

**Figure S16**. Correlations between the different experimental phases, separately for cognitive predictive learning (PL; S4) and fear learning (FL; studies S1, S2, S3, S5 and S6) paradigms (coloured lines) and for all studies combined (black line).

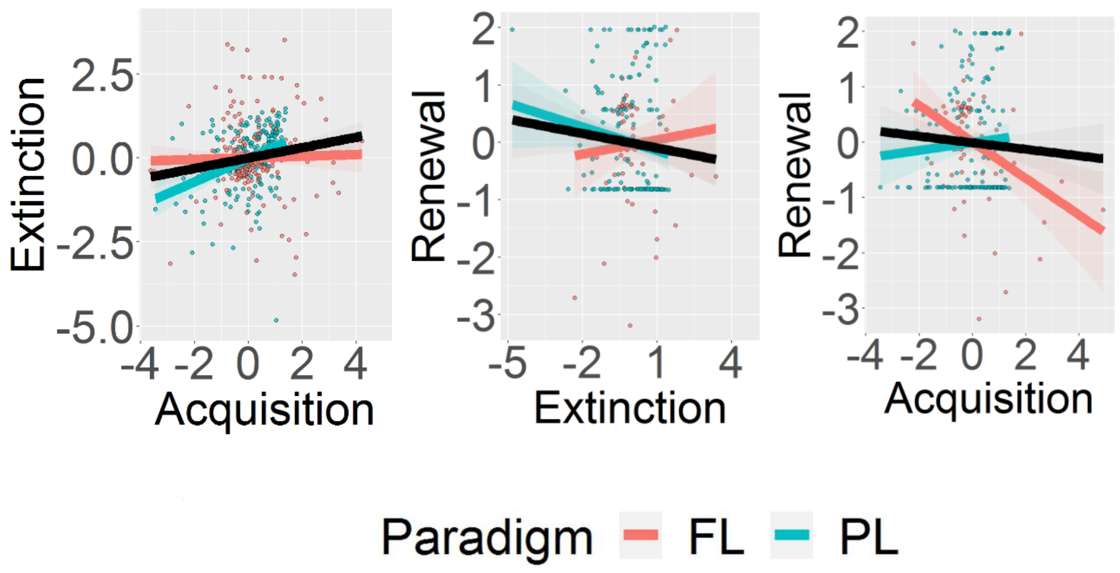

**Figure S17**. Multicollinearity diagnostics (Tolerance and VIF, upper left; Variance Proportion, upper right; Condition Index and Eigenvalue, bottom), showing there was low multicollinearity in our study.

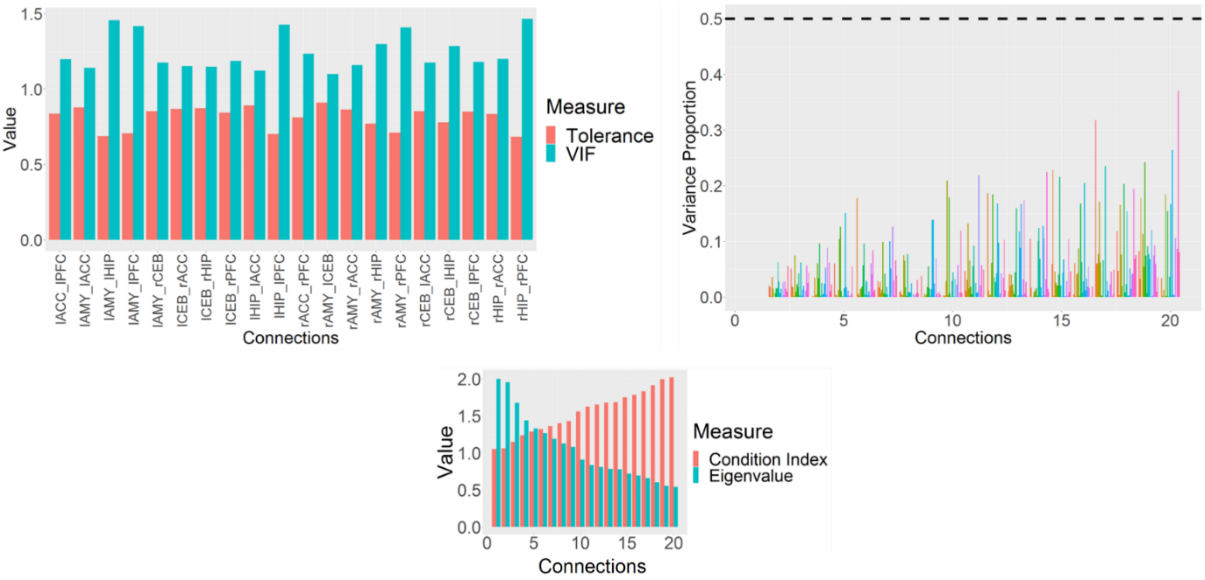

**Figure S18**. Correlations between Lasso coefficients and those obtained by standard ordinary least squares regression, Ridge regression and Elastic Net.

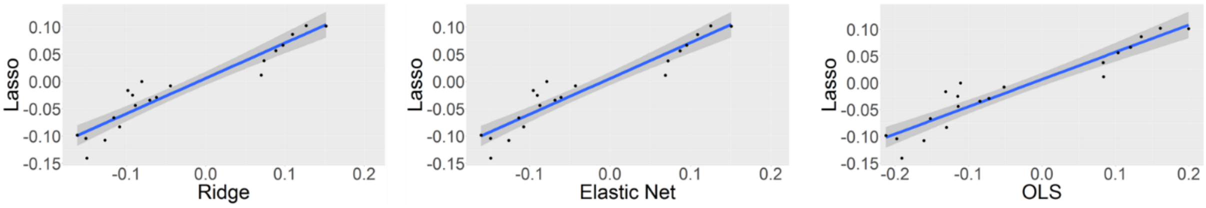

**Figure S19**. Histogram showing that the learning variable was normally distributed

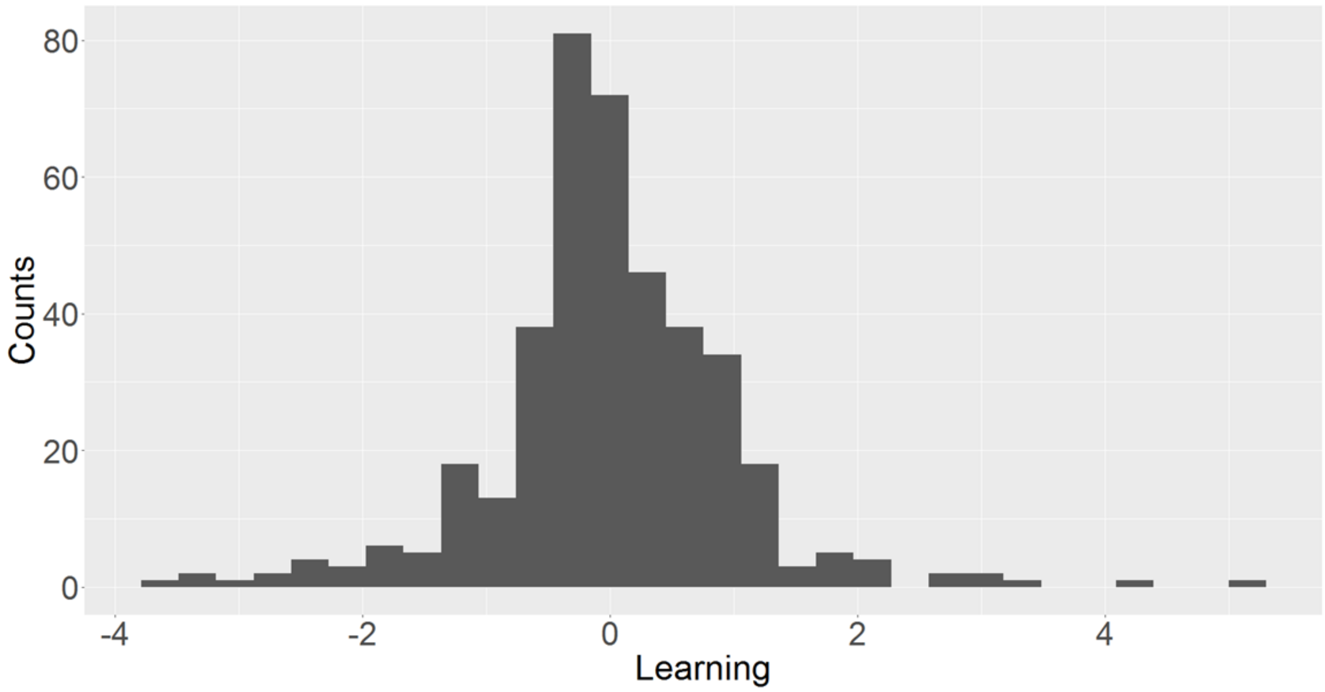

**Figure S20**. Simulation analysis using sample sizes of 90, 300, or 1000 observations. FL=Fear Learning studies (S1,S2,S3,S5,S6); PL=Predictive Learning studies (S4); FLr=Fear Learning renewal study (S2); PLr=Predictive learning renewal study (S4).

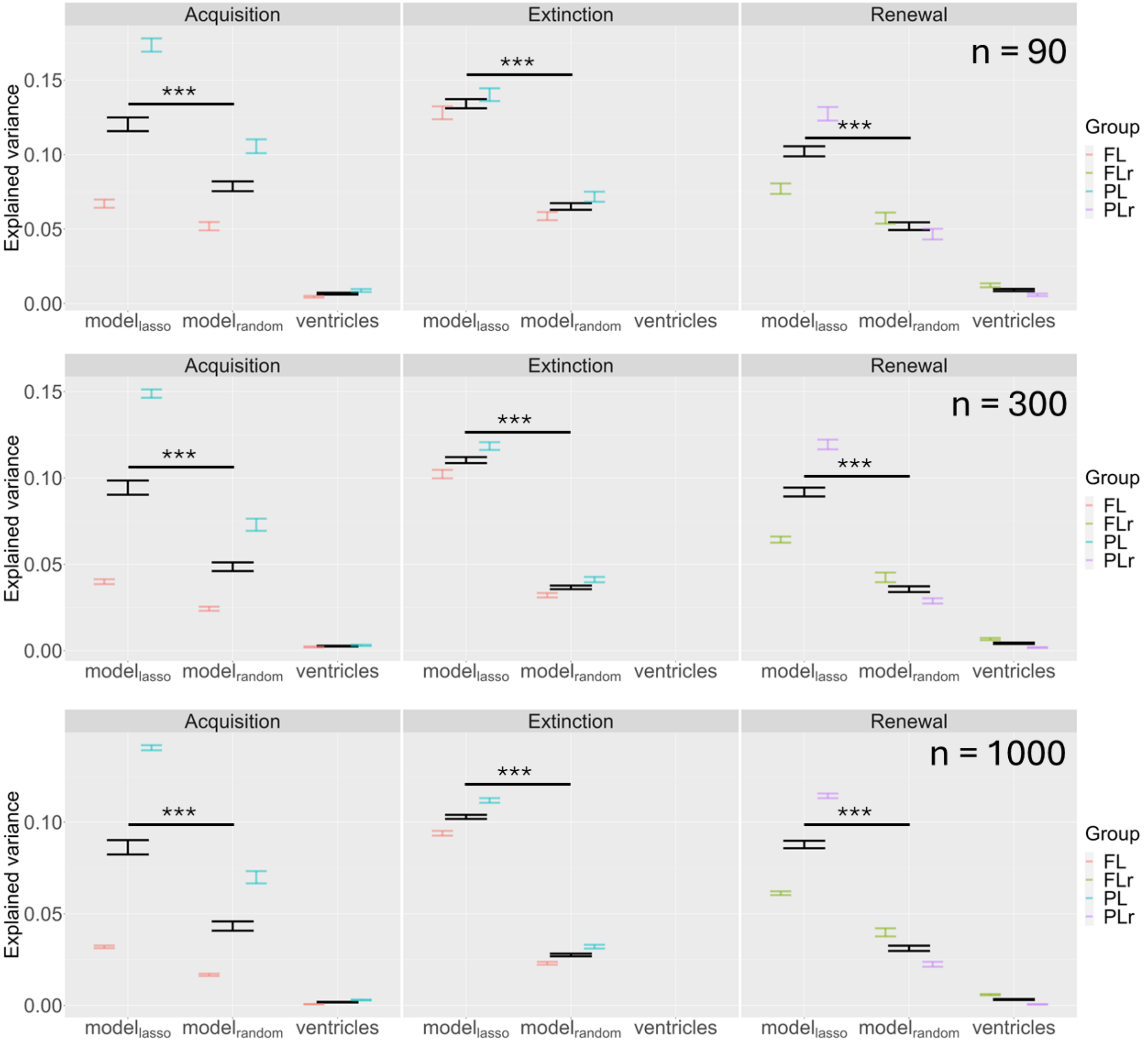

**Figure S21**. Average SCR learning estimates for CS+ and CS- for each phase and SCR study using electrical shocks as US (and same non-linear modelling strategy; see main text).

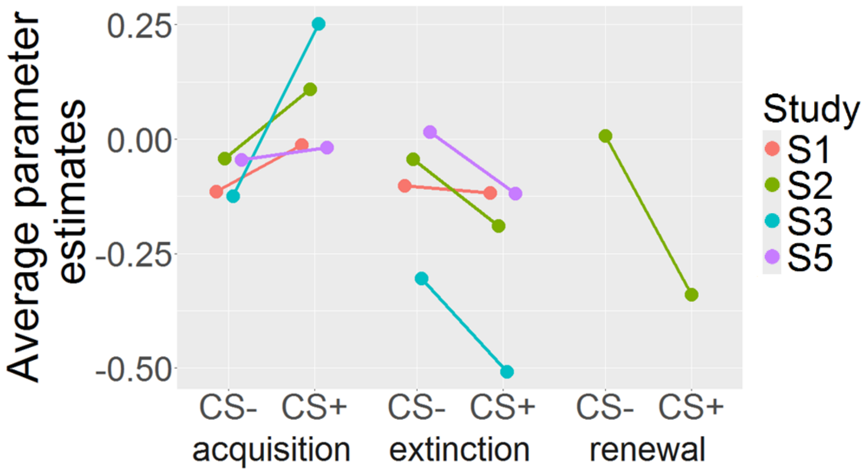

**Figure S22**. Significance of each predictor in the lasso model using traditional p-values. Stars indicate that the connection is selected significantly more often than expected by chance (Binomial test, p-value < 0.05, FDR adjusted for multiple comparisons).

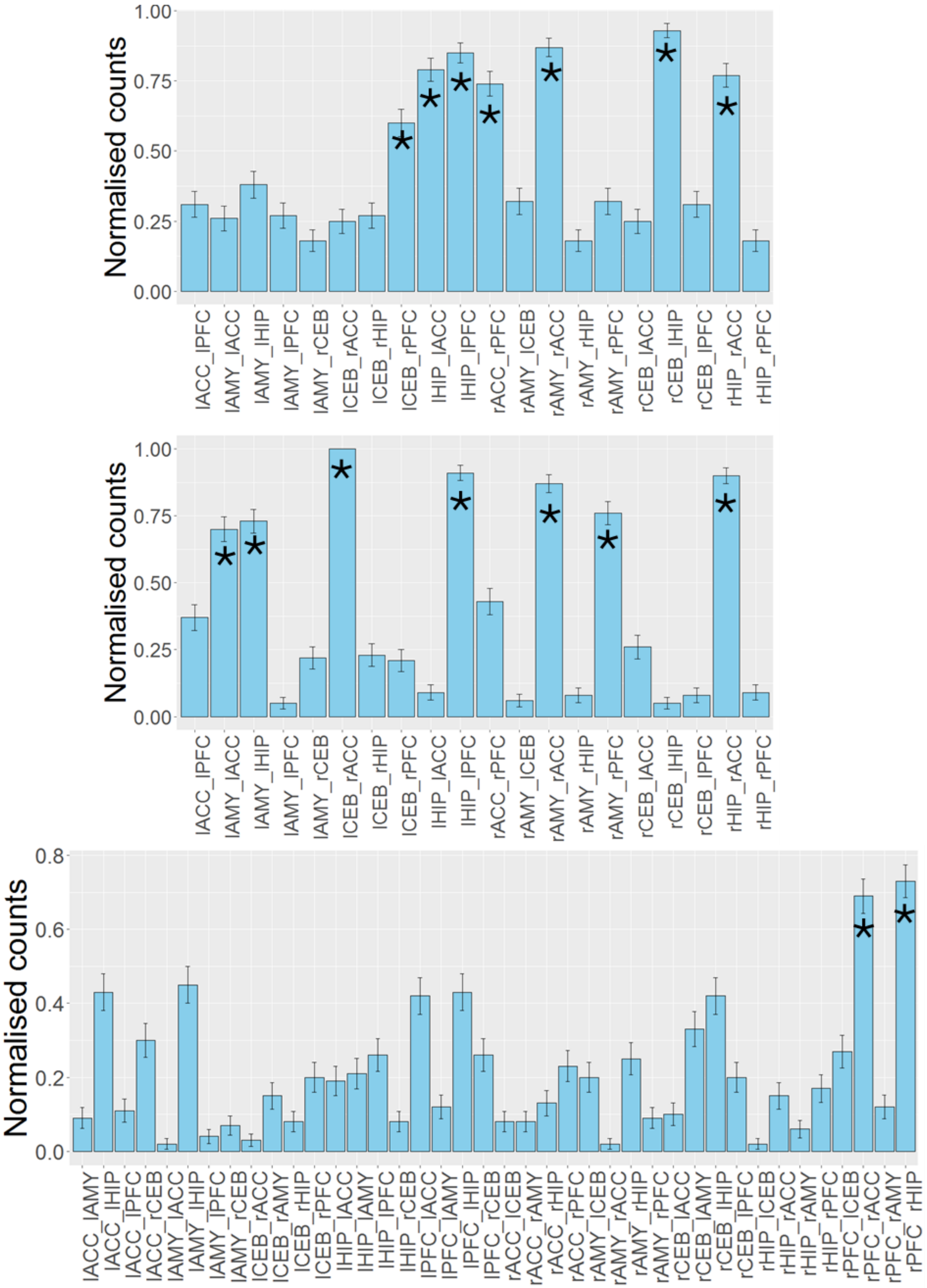

**Figure S23**. Regression models for all connections shown in Fig. 3B (acquisition). FC = Functional connectivity; SC = Structural connectivity; EC = Effective connectivity. All=All studies (S1,S2,S3,S4,S5,S6); FL=Fear Learning studies (S1,S2,S3,S5,S6); FLc=Fear Learning classical paradigm (S2,S3); FLs=Fear Learning similar paradigm (S2,S3,S5); PL=Predictive Learning studies (S4); FLst=Fear Learning studies with standard paradigm (S1,S2,S3,S5,S6). ACC=Dorsal anterior cingulate cortex; AMY=Amygdala; CEB=Cerebellar nuclei; HIP=Hippocampus; PFC=Ventro-medial prefrontal cortex.

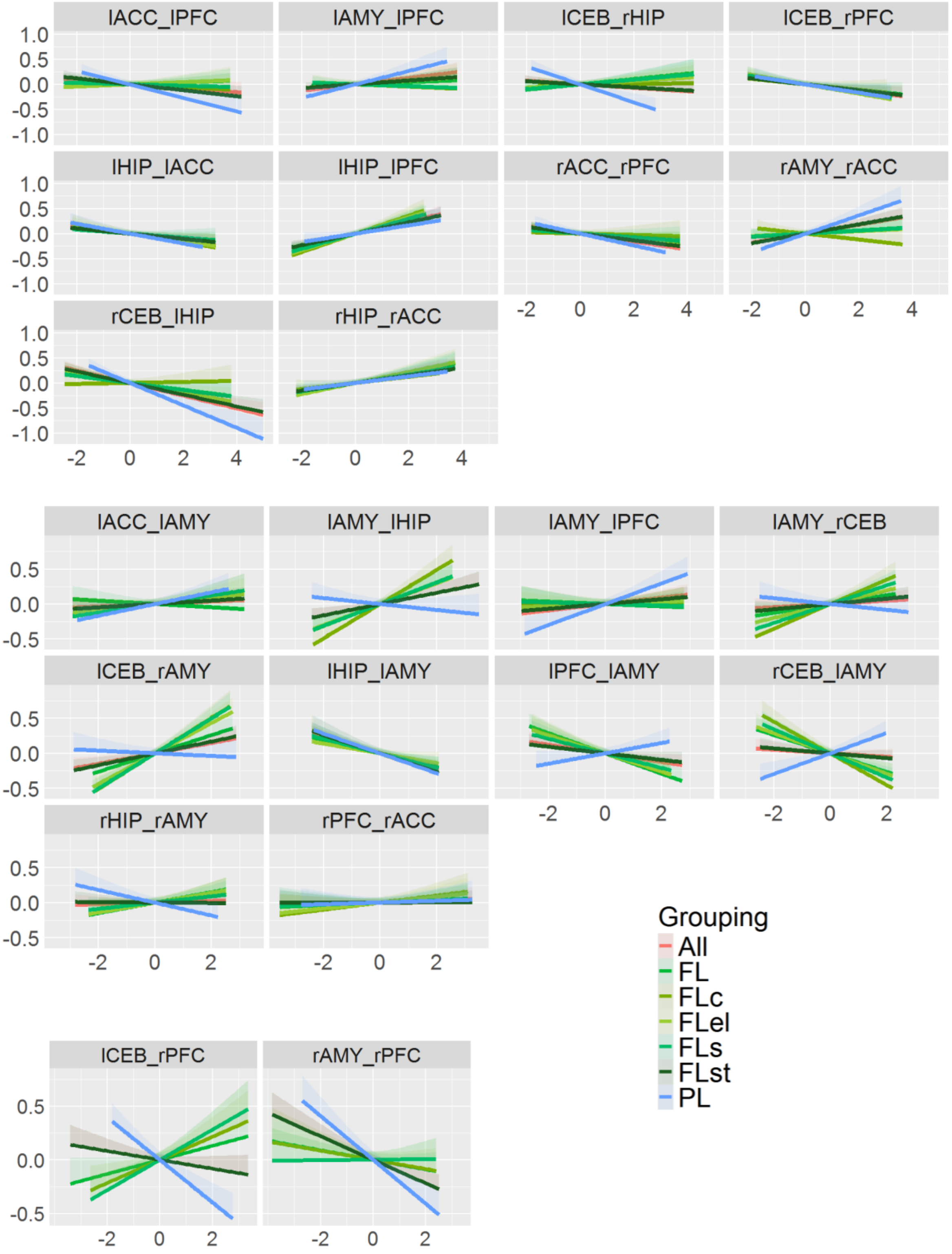

**Figure S24**. Regression models for all connections shown in Fig. 3C (extinction). Functional connectivity; SC = Structural connectivity; EC = Effective connectivity. All=All studies (S1,S2,S3,S4,S5,S6); FL=Fear Learning studies (S1,S2,S3,S5,S6); FLc=Fear Learning classical paradigm (S2,S3); FLs=Fear Learning similar paradigm (S2,S3,S5); PL=Predictive Learning studies (S4); FLst=Fear Learning studies with standard paradigm (S1,S2,S3,S5,S6). ACC=Dorsal anterior cingulate cortex; AMY=Amygdala; CEB=Cerebellar nuclei; HIP=Hippocampus; PFC=Ventro-medial prefrontal cortex.

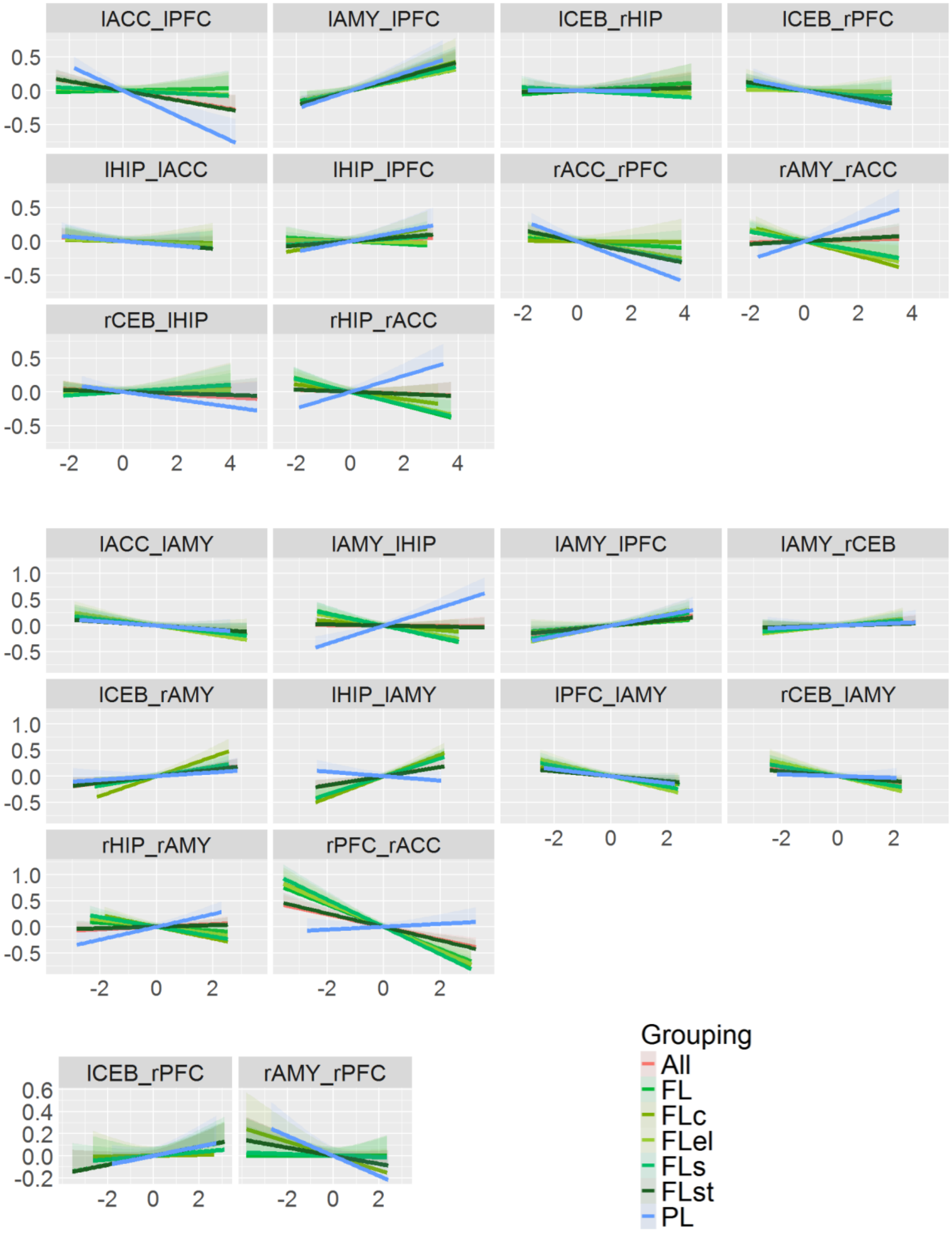

**Figure S25**. Regression models for all connections shown in Fig. 3C (renewal). Regression models for remaining connections not shown in Fig. 5E (right). EC = Effective connectivity. All=All studies (S1,S2,S3,S4,S5,S6); FL=Fear Learning studies (S2,S3,S5,S6); FLc=Fear Learning classical paradigm (S2,S3); FLs=Fear Learning similar paradigm (S2,S3,S5); PL=Predictive Learning studies (S1,S4); PLb=Predictive Learning behavioural studies (S4); SCR=SCR studies (S1,S2,S3,S5,S6). ACC=Dorsal anterior cingulate cortex; CEB=Cerebellar nuclei; HIP=Hippocampus; PFC=Ventro-medial prefrontal cortex.

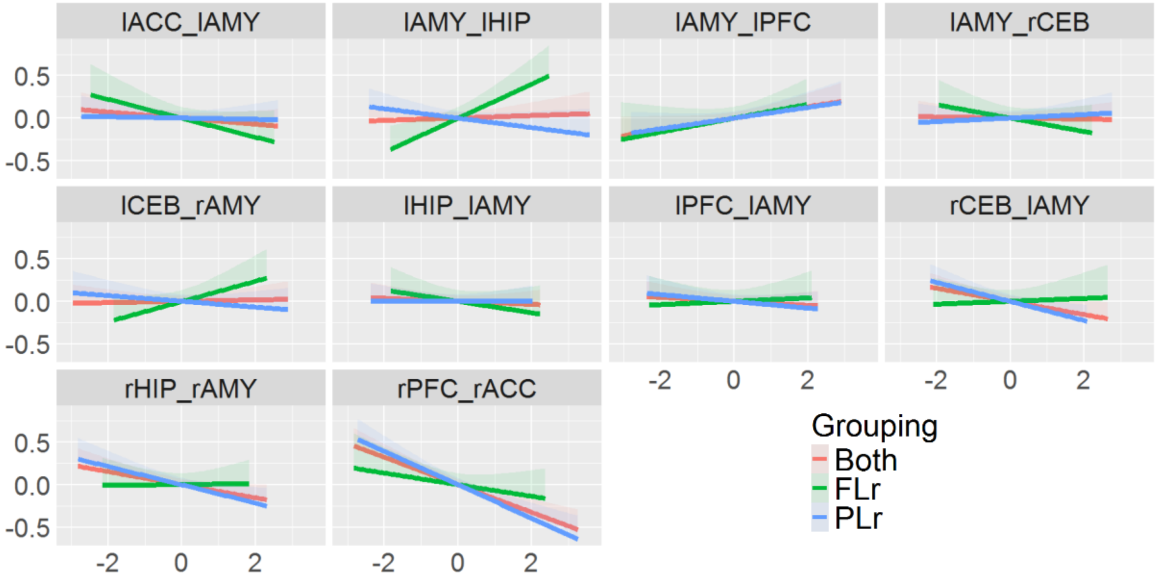

**Figure S26**. Learning was significant for all groupings for both acquistion (left) and extinction learning (right). FL=Fear Learning studies (S1,S2,S3,S5,S6); FLc=Fear Learning classical paradigm (S2,S3); FLel=Fear Learning electric shocks (S1,S2,S3,S5); FLr=Fear Learning renewal study (S2); FLs=Fear Learning similar paradigm (S2,S3,S5); FLst=Fear Learning standard paradigm (S2,S3,S5,S6).

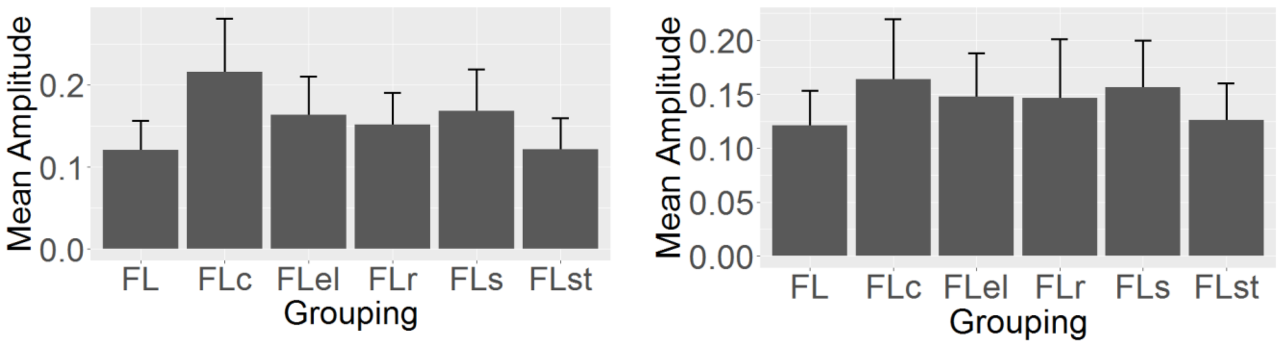

**Figure S27**. Significant connections for different fear learning groupings for acquisition. FC = Functional connectivity; SC = Structural connectivity; EC = Effective connectivity; FLst=Fear Learning standard paradigm (S2,S3,S5,S6); FLel=Fear Learning using electrical shocks (S1,S2,S3,S5); FLs=Fear Learning similar paradigm (S2,S3,S5). ACC=Dorsal anterior cingulate cortex; AMY=Amygdala; CEB=Cerebellar nuclei; HIP=Hippocampus; PFC=Ventro-medial prefrontal cortex.

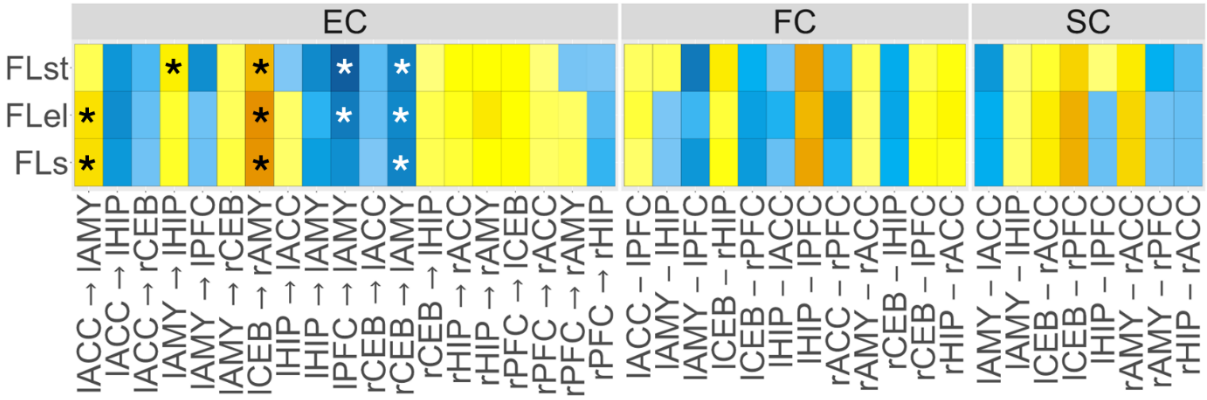

**Figure S28**. Significant connections for different fear learning groupings for extinction. FC = Functional connectivity; SC = Structural connectivity; EC = Effective connectivity; FLst=Fear Learning standard paradigm (S2,S3,S5,S6); FLel=Fear Learning using electrical shocks (S1,S2,S3,S5); FLs=Fear Learning similar paradigm (S2,S3,S5). ACC=Dorsal anterior cingulate cortex; AMY=Amygdala; CEB=Cerebellar nuclei; HIP=Hippocampus; PFC=Ventro-medial prefrontal cortex.

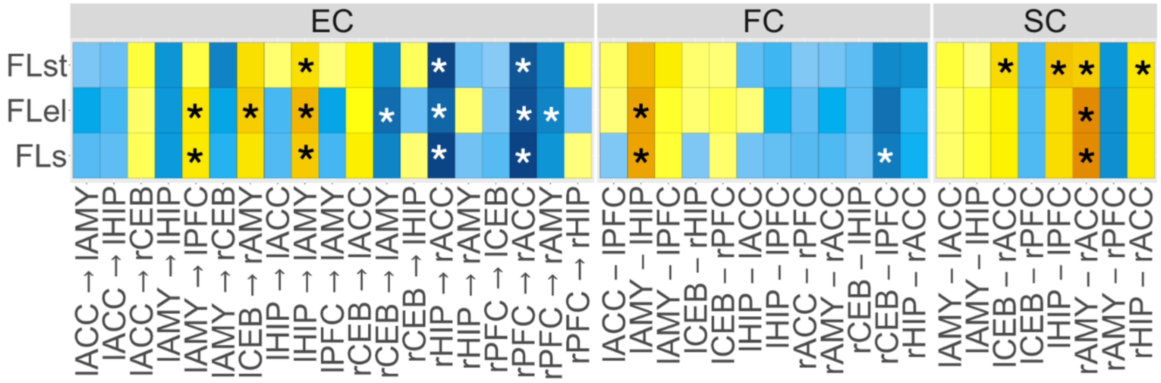

**Figure 29**: **Top:** Six resting-state networks [Schaeffer A. et al. (2018). Local-Global Parcellation of the Human Cerebral Cortex from Intrinsic Functional Connectivity MRI. Cerebral Cortex, 29:3095-3114] used in this analysis. **Middle**: Explained variance for each of the RSNs after re-running our entire FC pipeline. **Bottom**: Mean overlap between our fear and extinction network and the other RSNs. DMN=Default Mode Network (pink); EXN=Extinction Network; FPN=Frontoparietal Network (dark blue); LIN=Limbic Network (light blue); SAN=Salience Network (green); SMN=Somatomotor Network (red); DAN=Dorsal Attention Network (yellow).

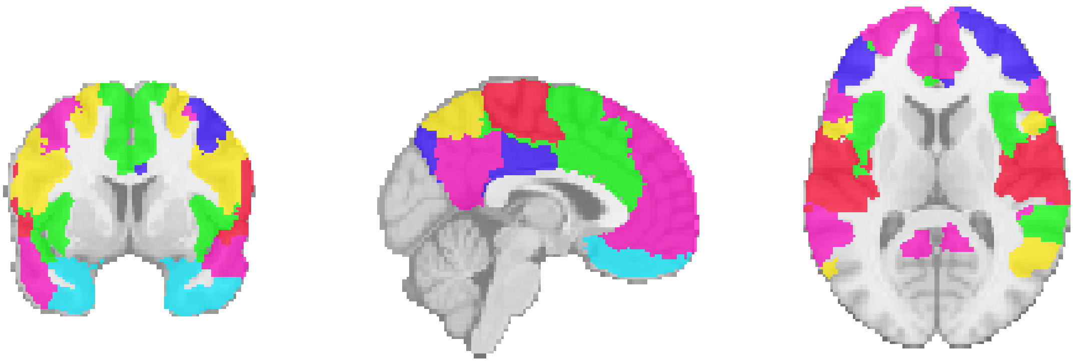

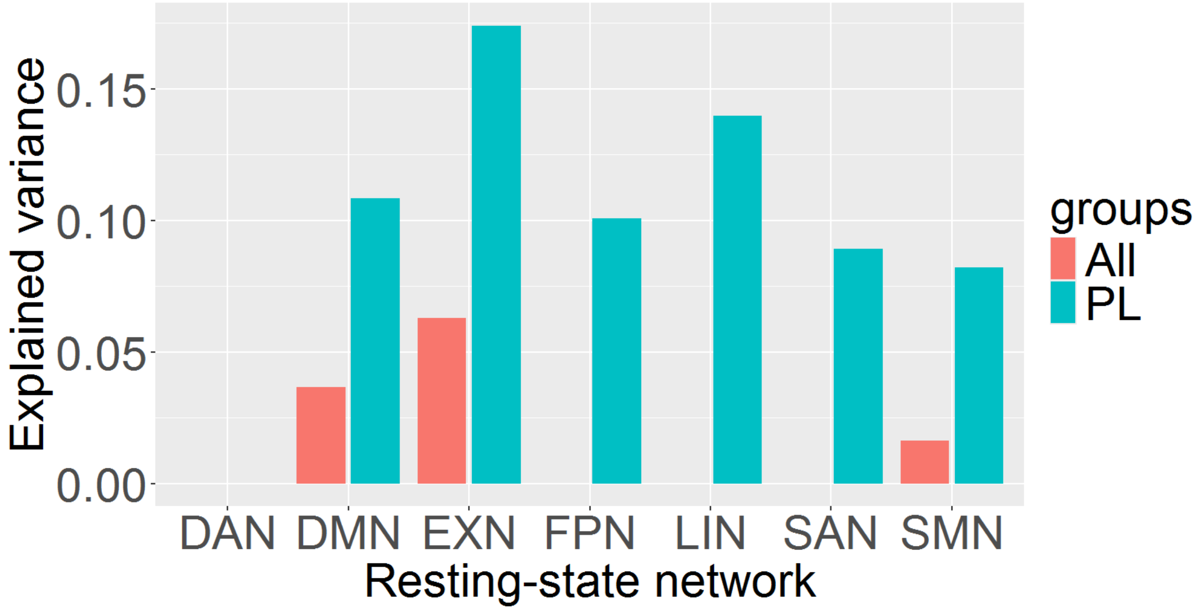


| RSN | Average number of overlapping voxels |
|---|---|
| LIN | 188 |
| DMN | 169 |
| SAN | 86 |
| FPN | 50 |
| DAN | 0 |
| SMN | 0 |

**Figure 30**. Test-retest reliability results for functional connectivity. Each colour represents a different session over the course of three days. The resting-state data analysed in the present study correspond to ses-01 run-1. The results for acquisition and extinction are shown on the top and bottom figures, respectively.

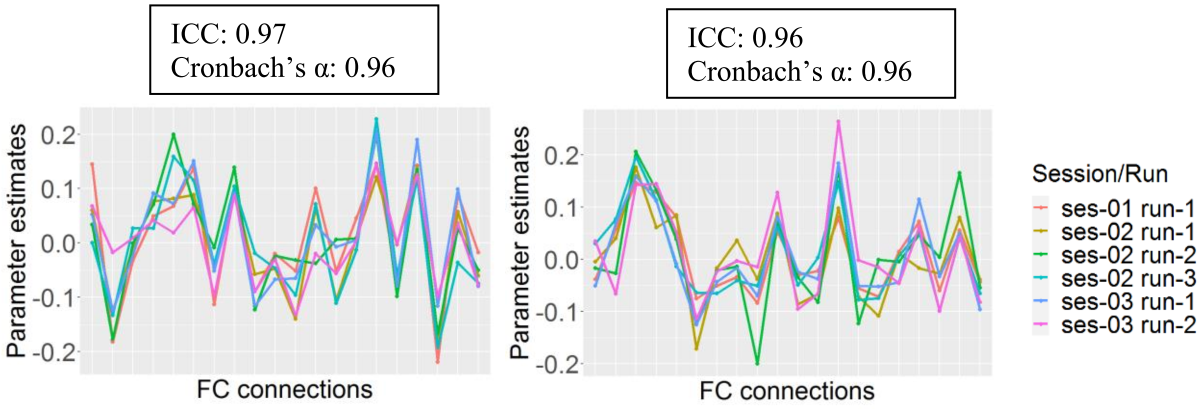

**Figure 31**. Test-retest reliability results for effective connectivity. Each colour represents a different session over the course of three days. The resting-state data analysed in the present study correspond to ses-01 run-1. The results for acquisition and extinction are shown on the top and bottom figures, respectively.

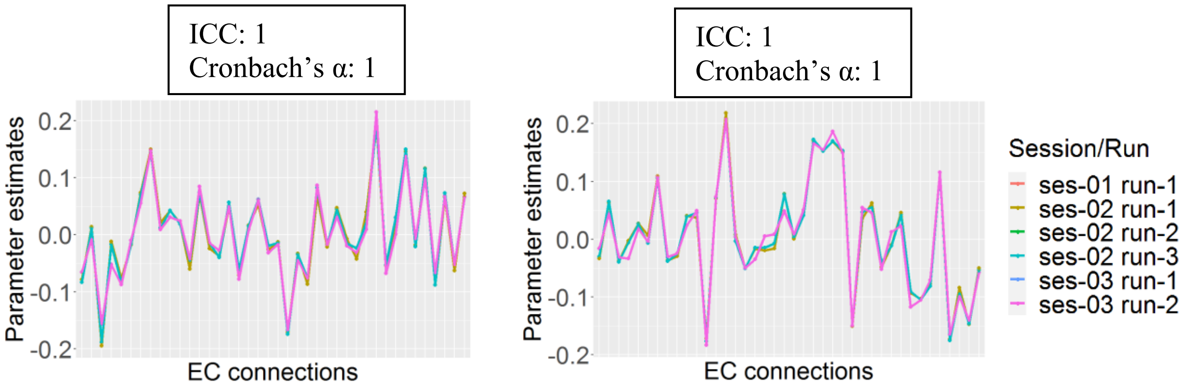

**Figure S32**. Generalisability results with non-overlapping train-test groups. FL=Fear Learning studies (S1,S2,S3,S5,S6); PL=Predictive Learning studies (S4).

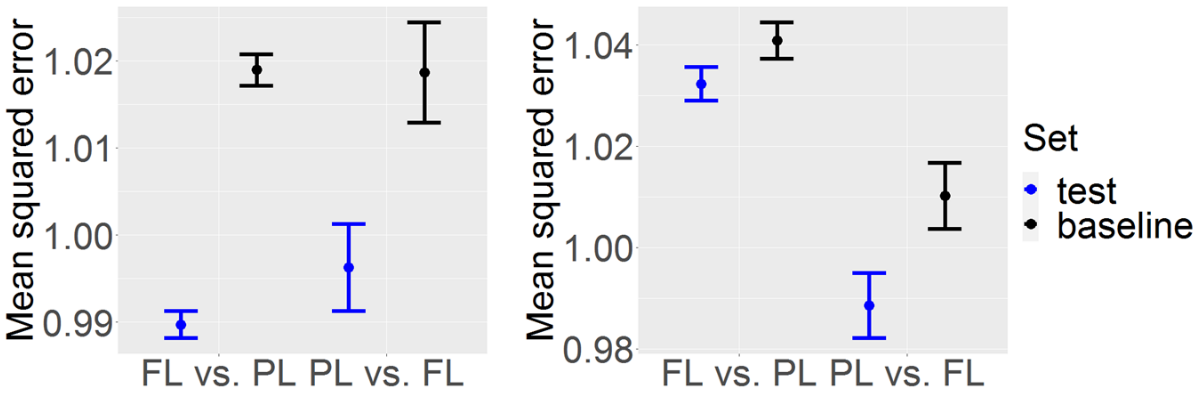

**Figure 33**. Short vs. long resting-state functional connectivity results. In all cases, correlations between datasets with different amount of volumes remained extremely high in all of the regions analysed in this study.

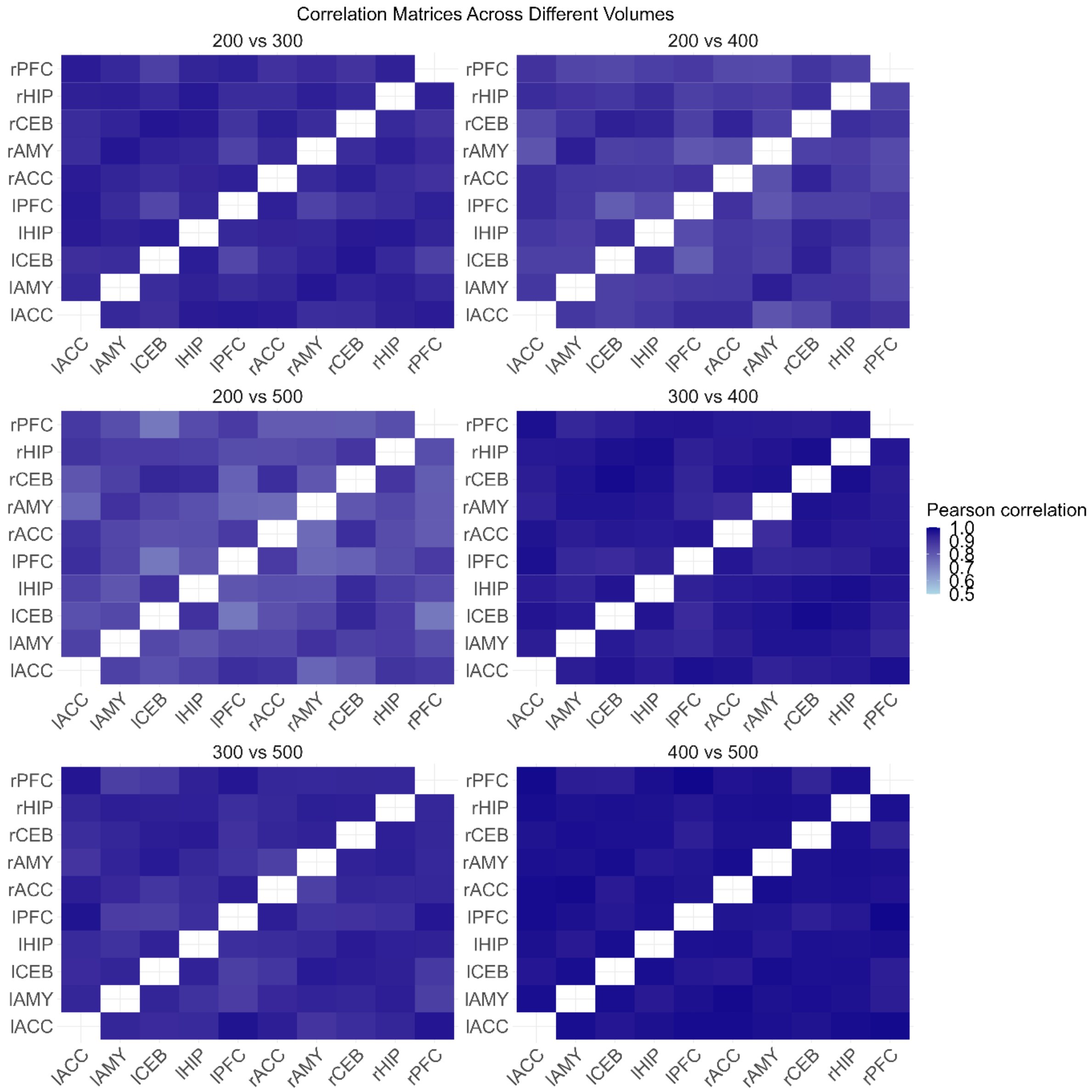

**Figure 34**. Ridge regression coefficients for rs-fMRI sessions with different durations. Although there were differences in parameter estimates among the different duration datasets, the relative differences between ROI pairs remained highly consistent across datasets with different amount of volumes.

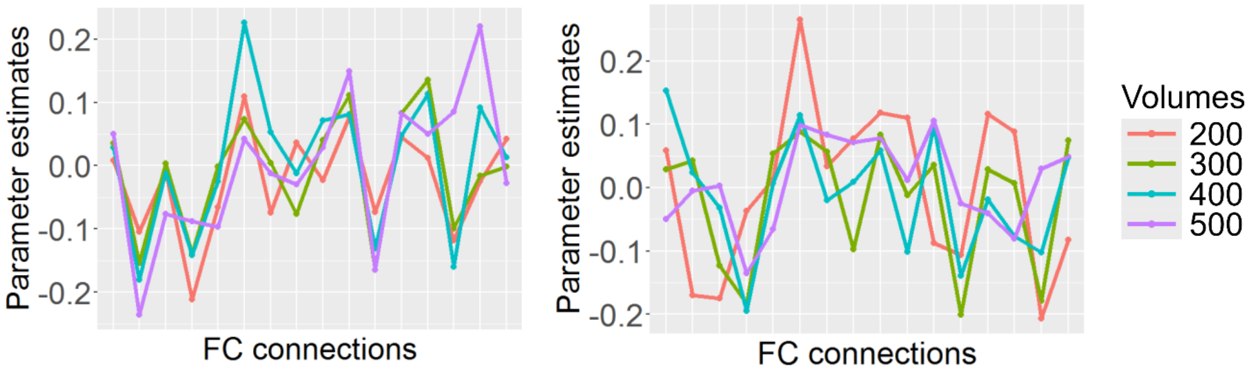

**Figure S35**. Simulation analysis using bootstrapping. FC = Functional connectivity; SC = Structural connectivity; EC = Effective connectivity.

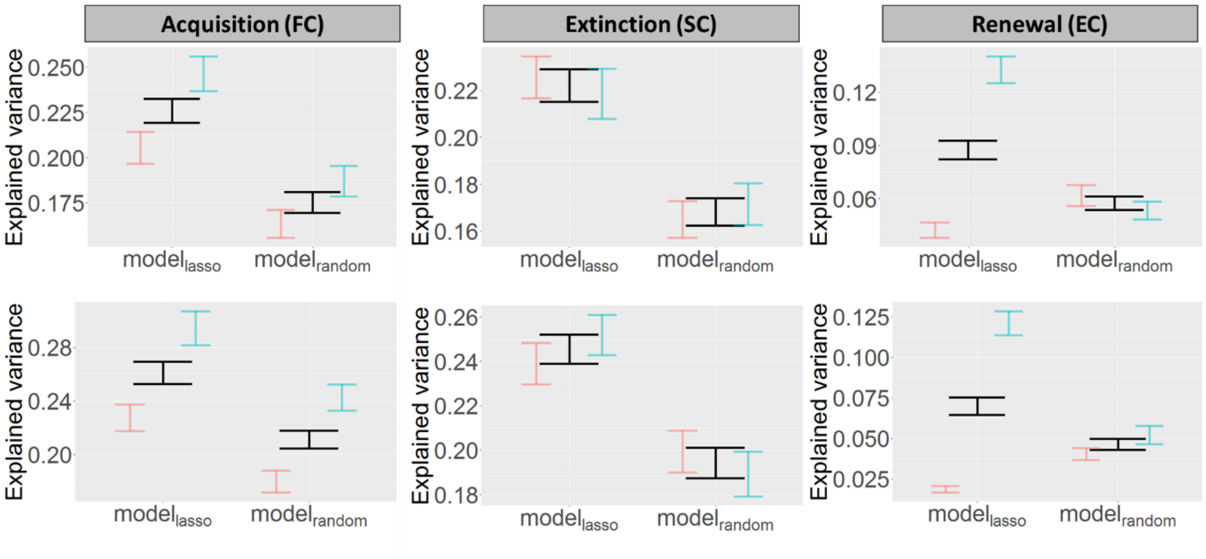

**Figure S36**. Activation foci of selected studies based on an extensive literature review of task-based fMRI studies. Dots indicate peak activations during acquisition (red) and extinction learning (blue) (see Table S1 for information on the selected studies).

**A.** Activation foci associated with acquisition and extinction learning

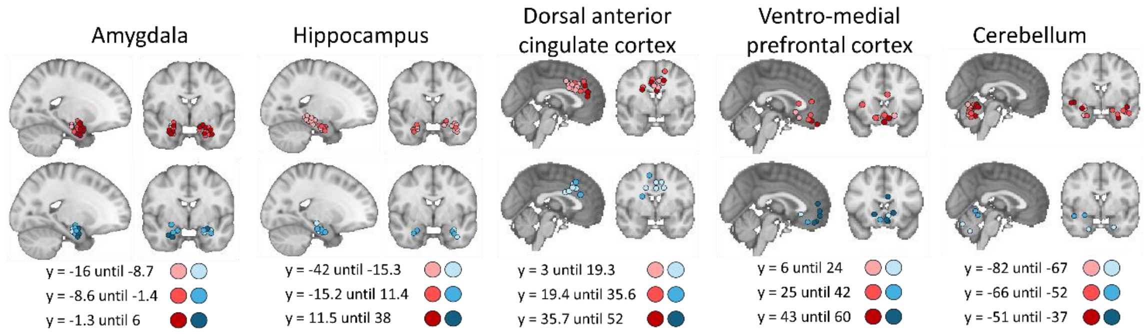

**Figure S37**. Correlation between the original behavioural (averaged) scores reported by Lissek et al. (2013; 2017; 2019) and the modelled responses as computed in our study. **Left**: Acquistion, $r = .94$, $p < .001$. **Right**: Extinction, $r = .98$, $p < .001$.

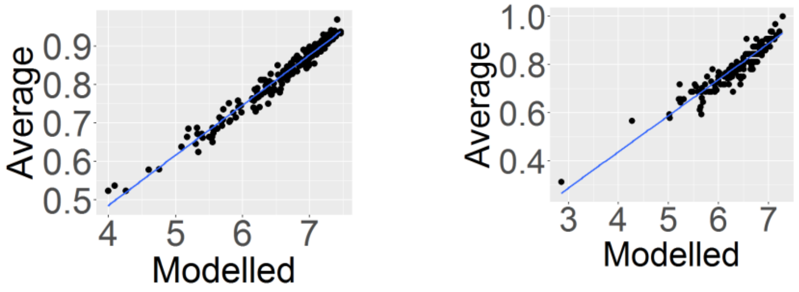
